# mRNA expression explains metabolic and thermal physiology

**DOI:** 10.1101/2022.01.19.477029

**Authors:** Melissa Drown, Douglas Crawford, Margie Oleksiak

## Abstract

Quantifying mRNA expression, which is heritable and physiologically inducible, reveals biologically important networks and pathways underlying complex traits. Here, we quantify mRNA expression in *Fundulus heteroclitus*, a small teleost fish, among three populations acclimated to 12°C and 28°C and relate it to variation in six, complex, physiological traits (whole animal metabolism (WAM), critical thermal maximum (CT_max_), and four substrate specific cardiac metabolic rates (CaM)). Although 366 heart mRNAs and 528 brain mRNAs had significant differential expression between the two acclimation temperatures, none of the mRNA acclimation responses were shared across all three populations in any tissue. Yet, within an acclimation temperature across all three populations, weighted gene co-expression network analyses show that mRNA expression patterns explained WAM, CT_max_, and CaM trait variation. These analyses revealed 9 significant heart MEs (first principal component of module expression) and 4 significant brain MEs. Heart MEs explained variation in WAM, CT_max_, and two of the four substrate specific cardiac metabolic rates at 12°C, and CT_max_ at 28C. In contrast, brain MEs explained CT_max_ and WAM at 28°C but not at 12°C. Combining MEs as multiple correlations, 82% of variation in WAM at 12°C was explained by four heart MEs, 80% of variation in fatty-acid CaM at 12°C was explained by three heart MEs, and 72% of variation in CT_max_ at 28°C was explained by three brain MEs. These MEs were enriched for Kyoto Encyclopedia of Genes and Genomes (KEGG) terms related to specific metabolic pathways, suggesting that they represent biologically relevant pathways. Together these data suggest that mRNA co-expression explains complex traits; moreover, physiological traits are more reliant on heart expression at 12°C and brain expression at 28°C.

**Author Summary:** Despite an abundance of genomic data, the molecular and genetic underpinnings of complex traits remain poorly understood. To better understand the molecular basis of complex traits, we used heart and brain mRNA expression to explain complex traits- physiological responses to temperature- in individuals collected from three saltmarsh fish (*Fundulus heteroclitus*) populations acclimated to 12°C and 28°C. We found that while physiological traits did not differ among populations, the mRNAs important for acclimation responses were >88% unique to a single population and differed between heart and brain tissues. We also found tissue specific co-expressed mRNAs that explain up to 82% of complex traits including whole animal metabolism, upper thermal tolerance, and substrate specific cardiac metabolism measured at 12°C or 28°C acclimation conditions. Notably, sets of co-expressed mRNAs related to these traits are enriched for molecular pathways affecting metabolism, giving insight into the molecular underpinnings of these traits.

## Introduction

Many genotype to phenotype mapping approaches such as expression quantitative trait loci (eQTL) mapping, co-expression network analysis, and functional genomics investigations (i.e., systems genetics) combine data across biological organization levels among inbred lines and leverage significant genomic resources (*e.g.,* well annotated genomes and transcriptomes, inbred lines, gene editing) (1). In several model organisms, these approaches have improved our ability to dissect complex trait architecture. For example, in fruit flies (*Drosophila*) variation in starvation resistance and startle response is explained by mRNA expression and polymorphic loci (2). In *Caenorhabditis elegans*, 199 recombinant inbred lines recently were used to identify 36 loci related to metabolism (3). In *Arabidopsis*, both segregating and isogenic lines were used to uncover complex genetic architecture of growth and morphology related traits (4).

In comparison to inbred lines, non-model organisms have more polymorphic alleles and greater heterozygosity, and thus non-model organism studies can provide a more nuanced understanding of how natural genetic variation impacts complex phenotypes (5, 6). Furthermore, non-model organism studies have become even more relevant with increased genomic resource availability and development of high-throughput approaches to measure behavioral and physiological traits under realistic ecological conditions (7–10).

Despite these recent advances, the genomic basis of well-studied traits, like metabolism, remain poorly understood, especially among a diversity of species. This may be largely due to the highly polygenetic nature of complex traits like metabolism that results in smaller average effect sizes, thus making it more challenging to detect nucleotide variation associated with physiological trait variation. Additionally, identifying genotype-to-phenotype associations is even more challenging when the genetic architecture is both polygenic and redundant because redundancy allows many small effect genetic polymorphisms associated with phenotypic variation to differ among environments, individuals, and populations (11–13). Although elucidating genotype-to-phenotype relationships is challenging for complex redundant polygenic traits, this knowledge is critical to understand evolutionary processes contributing to phenotypic variation.

One approach to the challenge of relating genotype to phenotype is to use mRNA expression, which provides three clear advantages. First, mRNA expression more readily explains trait variation than nucleotide variation (2, 6) because nucleotide variation is typically only binary while mRNA expression results from a combination of genetic polymorphisms.

These genetic polymorphisms collectively effect mRNA expression to provide a continuous trait distribution, and relating this polygenic continuous trait to more complex physiological traits is statistically stronger than relating binary nucleotide variation to complex traits. Second, quantifying mRNA expression is likely to provide more insight into the genetic architecture of physiological traits than genome wide association studies (GWAS) alone because mRNA expression provides information for both heritable and plastic responses (6, 14–19). That is, mRNAs capture multiple genetic effects including gene by environment interactions (GxE) in a single measure, which is essential when genetic architecture is context dependent (11, 13). Finally, a greater mechanistic understanding of complex traits is likely using mRNAs because mRNAs are more often defined genes associated with biochemical or physiological pathways (*e.g.,* through KEGG or Gene Ontology terms or molecular investigation). For example, mRNA expression variation across the tree of life explains phenotypes, including toxin response in yeast (20), flower induction in plants (21), diabetes (22), schizophrenia (23), cardiac metabolism (6), and many others (14, 24, 25). Overall, an improved mechanistic understanding of traits enables us to parse redundancy across biological organization levels and elucidate the evolutionary processes and genetic architectures contributing to phenotypic variation.

Here, we used mRNA expression patterns to identify molecular mechanisms underlying physiological phenotypes in the small saltmarsh teleost fish, *Fundulus heteroclitus,* captured from three wild populations and acclimated to 12°C and 28°C. To identify molecular mechanisms underlying physiological phenotypes, we quantified heart and brain mRNA expression among 86 *F. heteroclitus* individuals for which we had 6 measured physiological traits: whole animal metabolic rate (WAM), critical thermal maximum (CT_max_), and four substrate specific cardiac metabolic rates (CaM substrates: glucose [Glu], fatty acids [FA], lactate+ketones+ethanol [LKA], and endogenous ) (26). These traits are heritable and related to fitness, and CaM traits have been previously explained by mRNA expression variation (6, 27–31). In these fish, trait-specific acclimation responses to 12°C and 28°C differed. Specifically, acclimation responses eliminated temperature effects in CaM such that metabolic rates were the same when assayed at 12°C and 28°C, reduced the temperature effect in WAM to a small (1.2x) increase at higher temperatures (28°C), and enhanced CT_max_ by 6°C at higher temperatures (28°C) (26). Importantly, within an acclimation temperature individuals showed high trait variation with CVs (standard deviation/mean) for metabolic traits ranging from 22% to 55% (26). Presented here, we show that among these individuals, acclimation induced differential expression of 366 heart mRNAs and 528 brain mRNAs across all three populations, yet few differentially expressed mRNAs (one or less) were shared across all three populations. Within each acclimation temperature, co-expressed mRNA modules were significantly associated with WAM, CT_max_, and CaM. Using Kyoto Encyclopedia of Genes and Genomes (KEGG) and gene ontology (GO) enrichment, we identify biologically relevant networks among co-expressed mRNA modules that explain these traits. These data link a simpler molecular phenotype (mRNA expression) to complex trait variation to enhance our understanding of biological pathways that affect these traits and may be important for evolutionary adaptation.

## Results

*F. heteroclitus* used in this study were collected from three populations along the central coast of New Jersey, USA near the Oyster Creek Nuclear Generating Station (OCNGS), which produces a thermal effluent that locally heats the water. Three populations were sampled: 1) north reference (N.Ref; 39°52’28.000 N, 74°08’19.000 W), 2) south reference (S.Ref; 39°47’04.000 N, 74°11’07.000 W) and 3) a central site located between the southern and northern reference that is within the OCNGS thermal effluent (TE; 39°48’33.000 N, 74°10’51.000 W). The TE population used here differs by 4°C in habitat temperature from the two references populations (average summer high tide temperature 28°C N.Ref and S.Ref, and 32°C for TE) but is otherwise ecologically similar (26) with evidence that the TE population has locally adapted to life near the OCNGS (32). Prior to mRNA analyses for this study, individuals were acclimated to both 12°C and 28°C, and whole animal metabolic (WAM) and thermal tolerance traits (CTmax) were measured at both temperatures in all individuals. Next, substrate specific cardiac traits (CaM) were measured at either 12°C or 28°C (10). Finally, heart and brain samples were collected at the time of CaM measurements and prepared for mRNA sequencing. In total 219 individual hearts and brains were collected and used for mRNA sequencing.

### Sequencing analysis

mRNA expression was quantified by counting 3’ end reads from two Illumina HiSeq3000 lanes that yielded 10,535 mRNAs among hearts and 10,932 mRNAs among brains after filtering for 1.5 million reads per sample among hearts, 1 million reads per sample among brains, and at least 30 counts in 10% of individuals per mRNA with each tissue. In hearts this yielded an average of 8,224.7 reads per transcript and in brains an average of 6,578.5 reads per transcript. Sequencing statistics and sample sizes are summarized in Table 1. Using all heart and brain samples we examined mRNA expression variation using the top 500 most variable mRNAs for principal component analysis (PCA). While 86% of variance on PC1 clearly split 86 heart and brain samples into 2 distinct clusters, 25 samples (12 hearts, 13 brains) were clustered with the wrong tissue and were excluded (Fig. S1). The remaining 86 samples were used for all further tissue specific analyses. Using only these 86 individuals provides sufficient variation among individuals to examine the relationships between mRNA expression and physiology, although our analyses may be conservative because we removed 25 samples with ambiguous tissue expression.

**Table 1:** Sequencing statistics. Sequencing statistics and sample size distribution among tissues, acclimation temperatures, and populations.

| Heart Tissue | Sequencing Statistics |  | Acclimation Temperature | Population | Sample Size |
| --- | --- | --- | --- | --- | --- |
| Total N | 41 |  | 12°C | N.Ref | 4 |
| Total mRNAs | 10,535 |  |  | TE | 6 |
| Average reads per mRNA | 8,224.70 |  |  | S.Ref | 9 |
| Minimum reads per individual | 1.5 million |  | 28°C | N.Ref | 4 |
| Average reads per individual | 2.17 million |  |  | TE | 8 |
|  |  |  |  | S.Ref | 10 |
| Brain Tissue | Sequencing Statistics |  | Acclimation Temperature | Population | Sample Size |
| Total N | 45 |  | 12°C | N.Ref | 9 |
| Total mRNAs | 10,932 |  |  | TE | 11 |
| Average reads per mRNA | 6578.5 |  |  | S.Ref | 8 |
| Minimum reads per individual | 1 million |  | 28°C | N.Ref | 5 |
| Average reads per individual | 1.74 million |  |  | TE | 6 |
|  |  |  |  | S.Ref | 6 |

### Differential expression analysis

Differential expression patterns among populations and acclimation temperatures were identified using DESeq2 (33). First, to examine population and temperature specific expression we used model design: (∼Population + Acclimation-Temperature + Population*Acclimation-Temperature). This analysis revealed significant Population*Acclimation-Temperature interactions, suggesting acclimation temperature specific mRNA expression patterns among populations. Because of the significant interactions, we analyzed individuals acclimated to 12°C or 28°C separately with model design: ∼Population to identify differentially expressed mRNAs among populations within an acclimation temperature. Similar to other species, there were significant differentially expressed mRNAs between acclimation temperatures, reflecting changes in response to environmental temperature ((17, 34, 35), Table S1). Across all 3 populations, hearts had 366 mRNAs (3.5% of total) that were significantly different between the two acclimation temperatures (FDR <0.05) with equal up and down regulated for 12°C *versus* 28°C (183 up and 183 down). For brains, 528 mRNAs (4.8% of total) were significantly differently expressed between the two acclimation temperatures (FDR <0.05) with ∼2.5-fold more down regulated at 28°C relative to 12°C (148 up and 380 down).

While all three populations showed acclimation effects for heart and brain mRNAs, the affected mRNAs were not shared among all populations (Fig 1). For any single population, acclimation to 12°C and 28°C had significant mRNAs that were unique to each population (Table S1). For the 366 significant heart acclimation mRNAs, 94-98% were unique to a single population, and no acclimation heart mRNAs were shared across all three populations (Fig. 1A, C, Table S1). Similarly, for the 528 significant brain acclimation mRNAs, 88-96% were unique to one population (Fig. 1B, D, Table S1), and only one acclimated brain mRNA was shared across all populations. These heart and brain population specific mRNA acclimation responses were significant (chi-squared heart p=1.64×10^-10,^ brain p=2×10^-16^). Additionally, acclimation significant mRNAs were unique to either heart or brain with no shared (0 mRNAs) acclimation response between tissues. This reflected different expression patterns between tissues, previously identified with PCA analysis (Fig S1).

**Figure 1:**
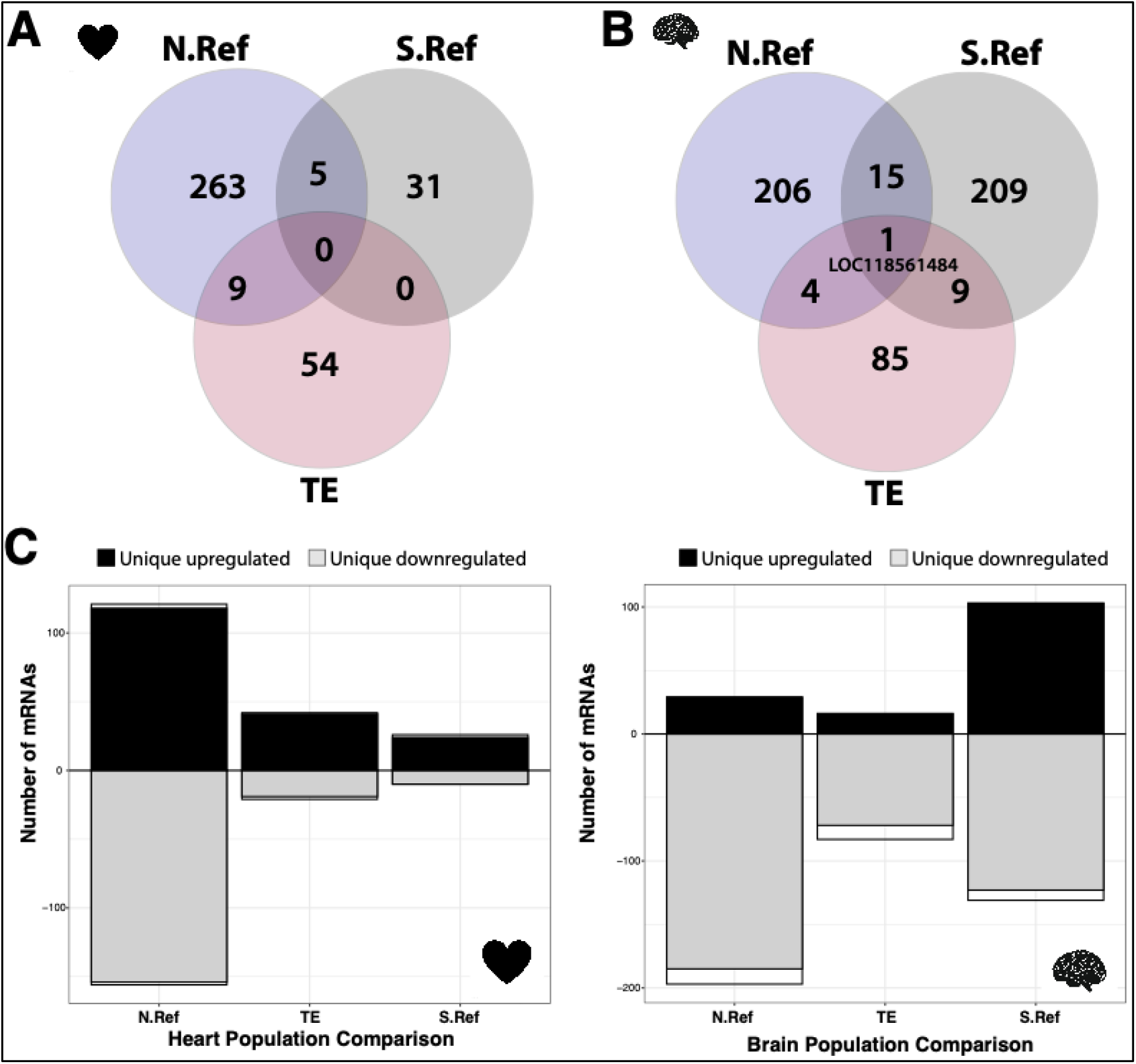
Population and tissue specific transcriptomic response to acclimation temperature. Number of differentially expressed mRNAs (DEGs) within each population between 12°C and 28°C acclimated hearts (**A and C**) and 12°C and 28°C acclimated brains (**B and D**). For both heart and brain, populations had many unique DEGs (**C and D**, upregulated=black, downregulated=grey) that were differentially expressed between acclimation temperatures and few shared DEGs (**C and D**, shared=white), with only 1 DEG shared among all three populations for brain (LOC118561484 in brains). Population and number of DEGs are not independent, Chi-Squared test, heart p=1.64×10^-10^, brain p=2×10^-16^.

At each acclimation temperature, populations also had significant expression differences (Fig. 2, Table S2). Hearts at 12°C and 28°C had 158 or 153 differentially expressed mRNAs among populations, respectively (Table S2). These represent 1.50% or 1.45% of all expressed heart mRNAs at 12°C and 28°C, respectively; brains had 242 or 330 differentially expressed mRNAs among populations at 12°C and 28°C, respectively. These represent 2.21% or 3.02% of all expressed brain mRNAs at 12°C and 28°C, respectively. None of the population effects were significant across all three populations (Fig. S3) for any acclimation temperature or tissue.

**Figure 2:**
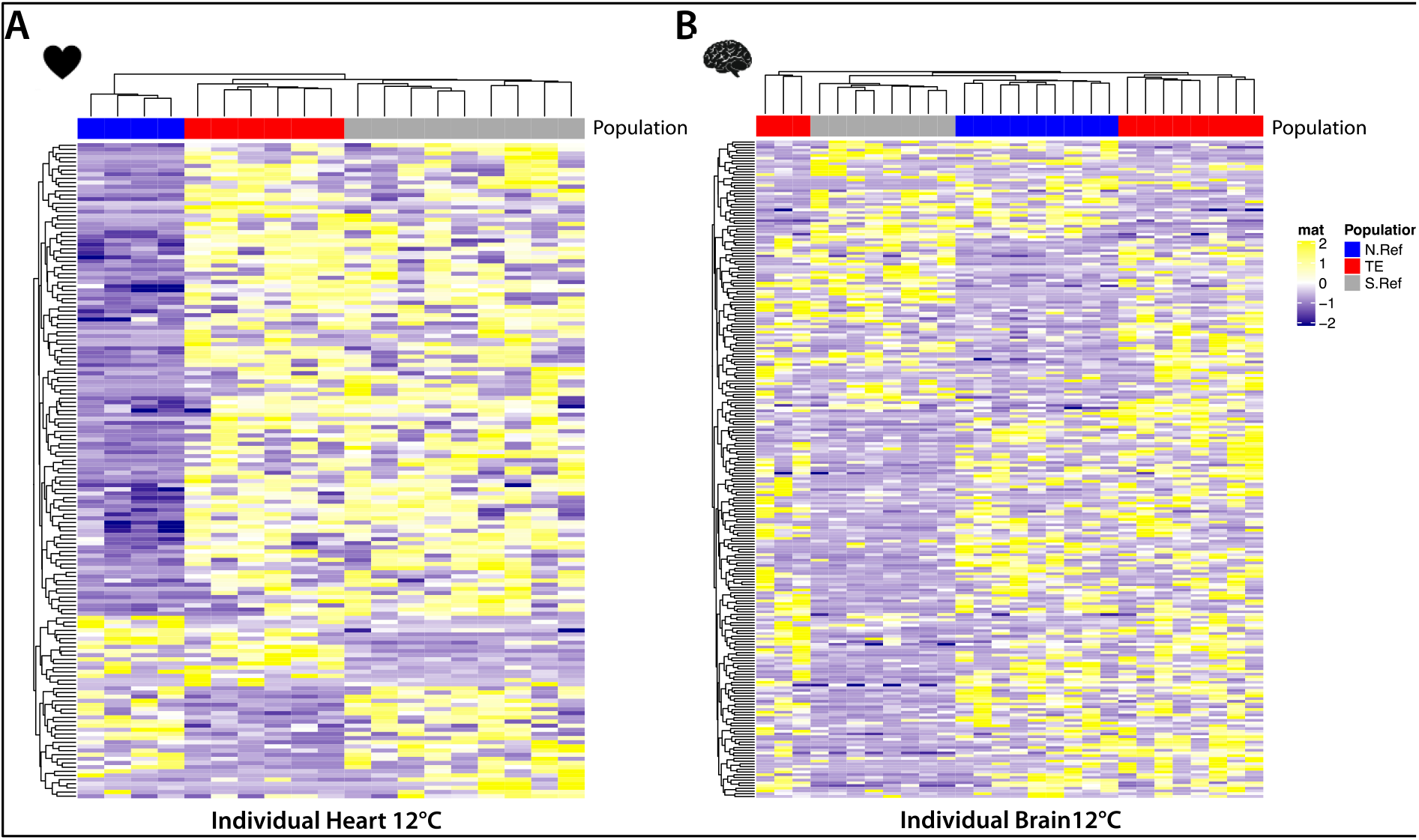
Temperature specific differential mRNA expression among populations. **A)** Differential expression for 158 heart mRNAs and **B)** 242 brain mRNAs between any two populations at 12°C. In hearts, 43.0% of mRNAs (68/158) are shared among any two population comparisons. In brains, 32.2% of mRNAs (78/242) are shared among any two population comparisons.

Importantly, there is an adaptive hypothesis: that the mRNAs in the anthropogenically warmed population, TE, are uniquely different, where TE is significantly different from both northern and southern reference populations with no significant differences between the references (17, 36). For heart mRNAs at 12°C there are 10 mRNAs (6.33% of significant mRNAs), and at 28°C there are 3 mRNAs (1.96%) where the TE population is uniquely different from both references (Fig S3, Table S2). For brain mRNAs at 12°C there are 11 mRNAs (4.55% of all significant mRNAs), and at 28°C there are 27 mRNAs (8.18%) where the TE population is uniquely different from both reference populations. While the overall frequency of differentially expressed genes is small (1.45% to 3.02% *vs.* total 10K mRNAs), the pattern where the TE population is different from both northern and southern reference populations but the two reference populations are not different is indicative of adaptation.

### Variation in mRNA expression

In addition to differential expression analysis, we were interested in the degree of mRNA expression variance. Previously, we found that variation in WAM, CT_max_, and substrate specific CaM was greater at 12°C than at 28°C. To determine if this was also true for mRNA expression, we quantified each mRNA’s coefficient of variation (CV, standard deviation/mean*100%) at 12°C and 28°C. For both heart and brain tissue there was greater average CV across all mRNAs at 12°C than at 28°C (T-test, heart p= 9.587e-05, brain p= 0.02014), similar to our findings of greater variation in physiological traits measured at 12°C.

### Weighted gene co-expression network analysis

We used weighted gene co-expression network analysis (WGCNA, (37)) to detect co-expressed mRNA clusters. WGCNA approaches group mRNAs with similar expression patterns into independent modules. Expression patterns for all mRNAs within a module were then summarized into principal components called module eigengenes (MEs, Table 2 for heart mRNAs and Table 4 for brain mRNAs), and these MEs were correlated to each of the six physiological traits (Table 3 shows significant heart MEs, and Table 5 shows significant brain MEs). Each ME has a “hub-MM”, the mRNA with the highest correlation to the ME, with MM being the correlation coefficient (Tables 2 and 4).

**Table 2:**
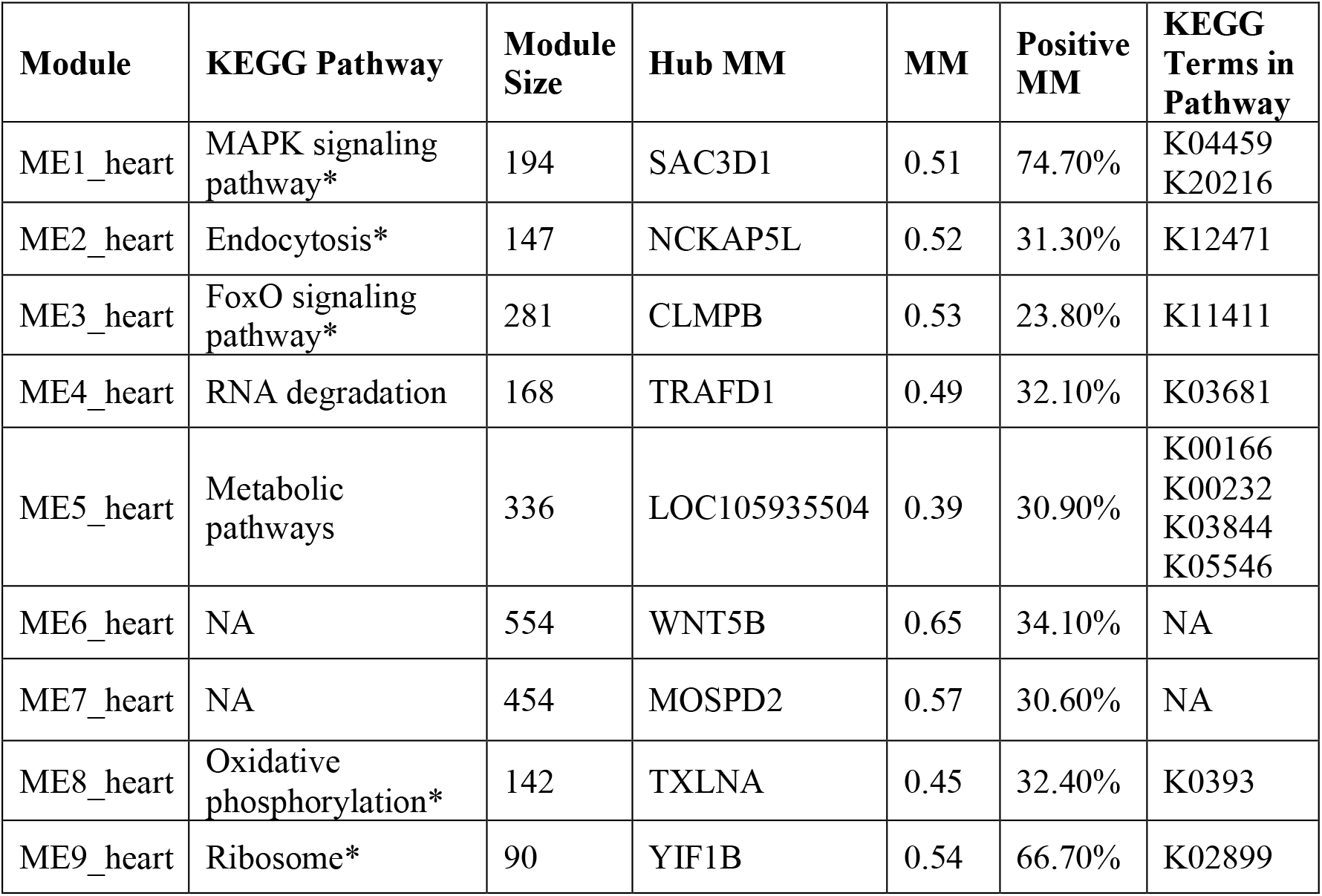
Heart Significant Modules. Columns include: Module = identifier for module eigengene (ME, first principal component of module), KEGG Pathway = top KEGG pathway determined by number of KEGG terms, Module Size = number of mRNAs in the module, Hub MM = mRNA with highest correlation with module eigengene, MM = correlation of hub mRNA with module eigengene, Positive MM = proportion of mRNAs with positive MM in the module, KEGG Terms in Pathway = enriched KEGG terms in the listed KEGG pathway. *Indicates modules where more than one KEGG Pathway had the same number of enriched KEGG terms, in which case the most informative KEGG pathway was selected.

**Table 3:** Heart Significant Module Trait Correlations. Significant heart ME correlations with FDR p<0.05. Columns include: Trait = traits significantly correlated with a given ME (critical thermal maximum: CT_max_, whole animal metabolic rate: WAM, cardiac metabolic rate: CaM with substrates fatty acids = FA, lactate+ketones+ethanol = LKA), Module = identifier for module eigengene (ME, first principal component of module), Correl coef = Pearson’s signed correlation coefficient for trait and ME, FDR P-value = multiple test corrected p-value for trait *versus* module correlation, Hub GS = mRNA with highest gene significance for the trait in the module, GS = gene significance, correlation between top module mRNA and trait, Positive GS = proportion of mRNAs in the module that are positively correlated with trait.

| Trait | Module | Correl coef | FDR P-value | Hub GS | GS | Positive GS |
| --- | --- | --- | --- | --- | --- | --- |
| CTMax 12°C | ME1_heart | 0.49 | 2.10E-02 | GPM6AA | -0.51 | 54.60% |
| CTMax 28°C | ME2_heart | -0.53 | 9.50E-03 | NCKAP5L | -0.50 | 35.30% |
| FA 12°C | ME3_heart | 0.50 | 1.02E-02 | PDCD4A | 0.55 | 65.80% |
| FA 12°C | ME4_heart | 0.53 | 4.82E-03 | RSRC1 | -0.58 | 13.10% |
| FA 12°C | ME5_heart | 0.55 | 4.82E-03 | RPL5A | 0.62 | 81.80% |
| heart mass 12°C | ME6_heart | -0.57 | 1.80E-03 | WRAP53 | 0.72 | 48.20% |
| LKA 12°C | ME6_heart | -0.65 | 3.39E-05 | LOC118563371 | 0.80 | 46.90% |
| LKA 12°C | ME7_heart | -0.56 | 1.29E-03 | LOC105934375 | 0.80 | 44.70% |
| WAM 12°C | ME8_heart | -0.56 | 1.20E-03 | LOC118563115 | 0.70 | 36.60% |
| WAM 12°C | ME9_heart | -0.55 | 1.20E-03 | LRRC58B | -0.56 | 10.00% |
| WAM 12°C | ME4_heart | -0.54 | 1.74E-03 | LOC118563570 | 0.61 | 65.50% |
| WAM 12°C | ME5_heart | -0.56 | 1.20E-03 | HPRT1L | 0.66 | 40.50% |
|  | AVERAGE | 0.55 |  |  |  |  |

**Table 4:** Brain Significant Modules. Columns include: Module = identifier for module eigengene (ME, first principal component of module), KEGG Pathway = top KEGG pathway determined by number of KEGG terms, Module Size = number of mRNAs in the module, Hub MM = mRNA with highest correlation with module eigengene, MM = correlation of hub mRNA with module eigengene, Positive MM = proportion of mRNAs with positive MM in the module, KEGG Terms in Pathway = enriched KEGG terms in the listed KEGG pathway.

| Module | KEGG Pathway | Module Size | Hub MM | MM | Positive MM | KEGG Terms in Pathway |
| --- | --- | --- | --- | --- | --- | --- |
| ME1_brain | Metabolic pathways | 393 | IDH3B | 0.47 | 26.40% | K00323<br>K01443<br>K10106<br>K19006 |
| ME2_brain | Tight junction | 198 | LOC105921153 | 0.52 | 28.20% | K07198<br>K08018<br>K08020 |
| ME3_brain | Lysosome | 342 | LRCH3 | 0.58 | 25.10% | K01134<br>K08568 |
| ME4_brain | RNA degradation* | 142 | SI:CH211-244C8.4 | 0.36 | 21.10% | K03681 |

**Table 5:**
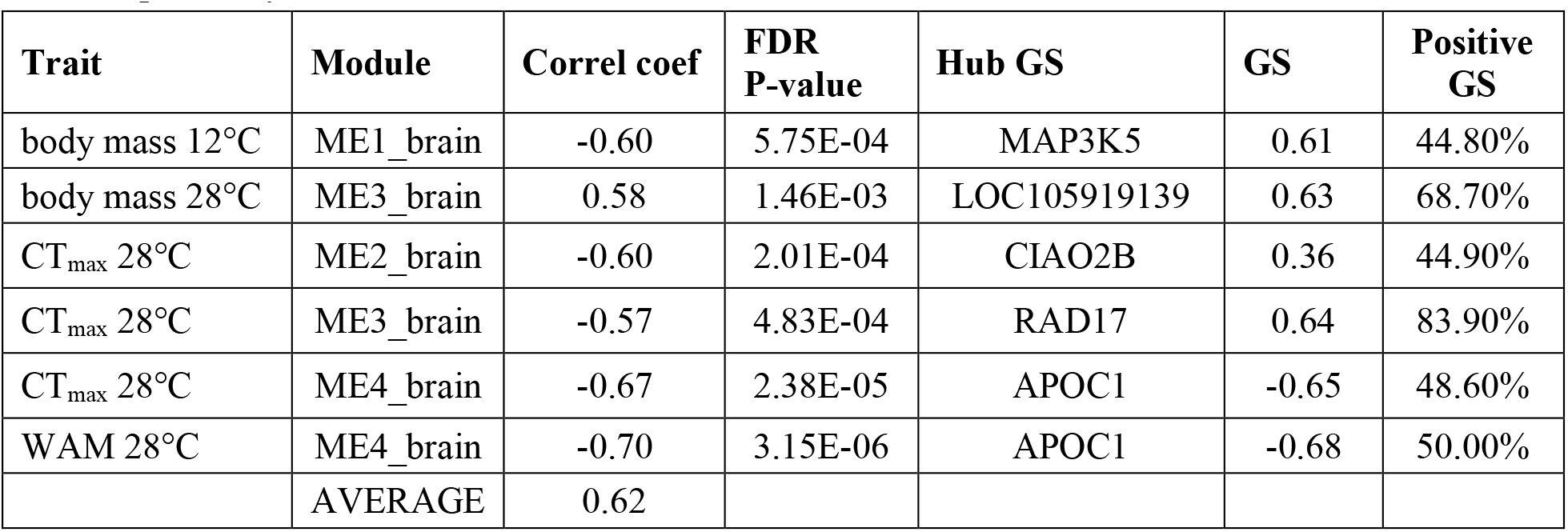
Brain Significant Module Trait Correlations. Significant brain *versus* ME correlations with FDR p<0.05. Columns include: Trait = traits significantly correlated with a given ME (critical thermal maximum: CT_max_, whole animal metabolic rate: WAM), Module = identifier for module eigengene (ME, first principal component of module), Correl coef = Pearson’s signed correlation coefficient for trait and ME, FDR P-value = multiple test corrected p-value for trait *versus* module correlation, Hub GS = mRNA with highest gene significance for the trait in the module, GS = gene significance, correlation between top module mRNA and trait, Positive GS = proportion of mRNAs in the module that are positively correlated with trait.

| Trait | Module | Correl coef | FDR P-value | Hub GS | GS | Positive GS |
| --- | --- | --- | --- | --- | --- | --- |
| body mass 12°C | ME1_brain | -0.60 | 5.75E-04 | MAP3K5 | 0.61 | 44.80% |
| body mass 28°C | ME3_brain | 0.58 | 1.46E-03 | LOC105919139 | 0.63 | 68.70% |
| $CT_{max}$ 28°C | ME2_brain | -0.60 | 2.01E-04 | CIAO2B | 0.36 | 44.90% |
| $CT_{max}$ 28°C | ME3_brain | -0.57 | 4.83E-04 | RAD17 | 0.64 | 83.90% |
| $CT_{max}$ 28°C | ME4_brain | -0.67 | 2.38E-05 | APOC1 | -0.65 | 48.60% |
| WAM 28°C | ME4_brain | -0.70 | 3.15E-06 | APOC1 | -0.68 | 50.00% |
|  | AVERAGE | 0.62 |  |  |  |  |

MEs were correlated to the body mass residuals for the six traits (WAM, CTmax, and the four substrate specific CaM). These analyses were done across all three populations because populations did not have any significant differences among traits. Each of the ME-trait correlations had a “hub-GS” – the mRNA in the module with the highest correlation to the trait, with GS (gene specific) being the correlation coefficient for this single mRNA. Both heart and brain WGCNA analysis used a minimum module size of 30 mRNAs per module and combined modules with correlation >75% (see methods). To verify that trait *versus* ME correlations were not driven by spurious outliers, we used a jack-knife approach to subsample 90% of individuals and repeat ME-trait correlations 100 times. Correlations that were significant in at least 70 out of 100 repeated correlations in the same direction (positive or negative correlation coefficient) were retained for further analysis (see methods).

#### Heart WGCNA

For heart mRNAs we found 39 co-expression modules with 90 to 554 mRNAs in each module (Table 2), and these heart MEs (first principal component of co-expressed mRNAs) were correlated to six physiological traits at each acclimation temperature. There were 12 significant ME-trait correlations: 9 heart MEs with 5 temperature specific traits (FDR <0.05, Table 3, Fig 3 and 4). Traits correlated with at least one of these 9 heart MEs included: at 12°C WAM, CT_max_, FA CaM, heart mass, LKA CaM, and at 28°C, CTmax (Table 3, Fig. 3, 4). Two of these modules (ME4_heart, ME5_heart) were correlated with both WAM at 12°C and FA CaM at 12°C, and one module (ME6_heart) was correlated with both LKA CaM at 12°C and heart mass at 12°C. WAM at 12°C had the most significant ME correlations (4 total), followed by FA CaM at 12°C (3 total) and LKA CaM at 12°C (2 total). The other three traits were each correlated with a single module (Table 3). On average, a single heart module explained 55% of variance for one trait with a minimum of 48.5% (ME1_heart with CT_max_ 12°C) and a maximum of 65% (ME6_heart with LKA 12°C).

**Figure 3:**
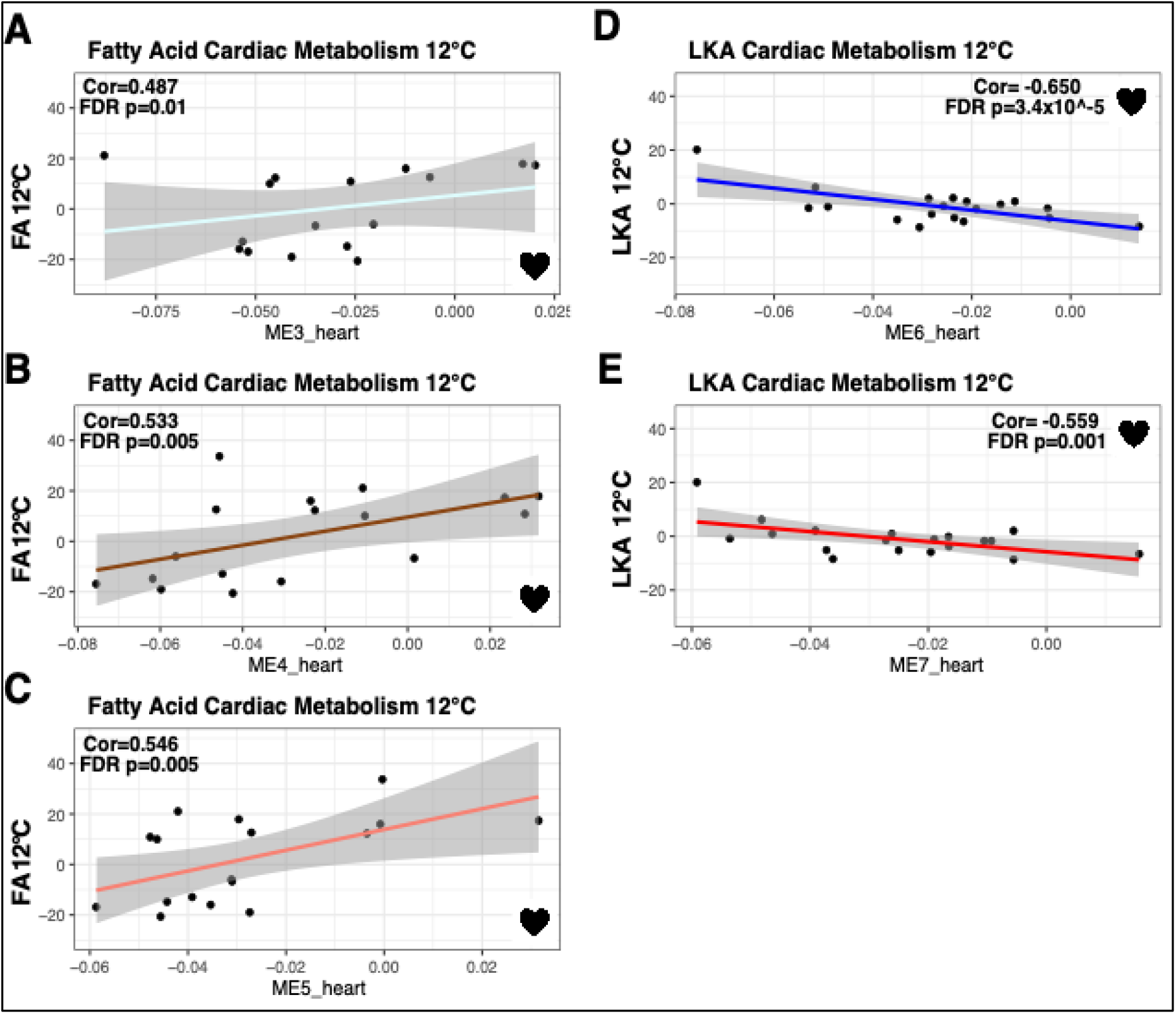
Significant cardiac metabolism-heart module correlations from weighted gene co-expression network analysis. Significant correlation of fatty acid cardiac metabolic rate at 12°C (N=16) with ME3_heart (**A**), ME4_heart (**B**), and ME5_heart (**C**). Significant correlation of lactate, ketone, and alcohol (LKA) cardiac metabolic rate at 12°C (N=19) with ME6_heart (**D**) and ME7_heart (**E**). Pearson correlation coefficients (Cor) and FDR p-values are displayed for each significant correlation.

**Figure 4:**
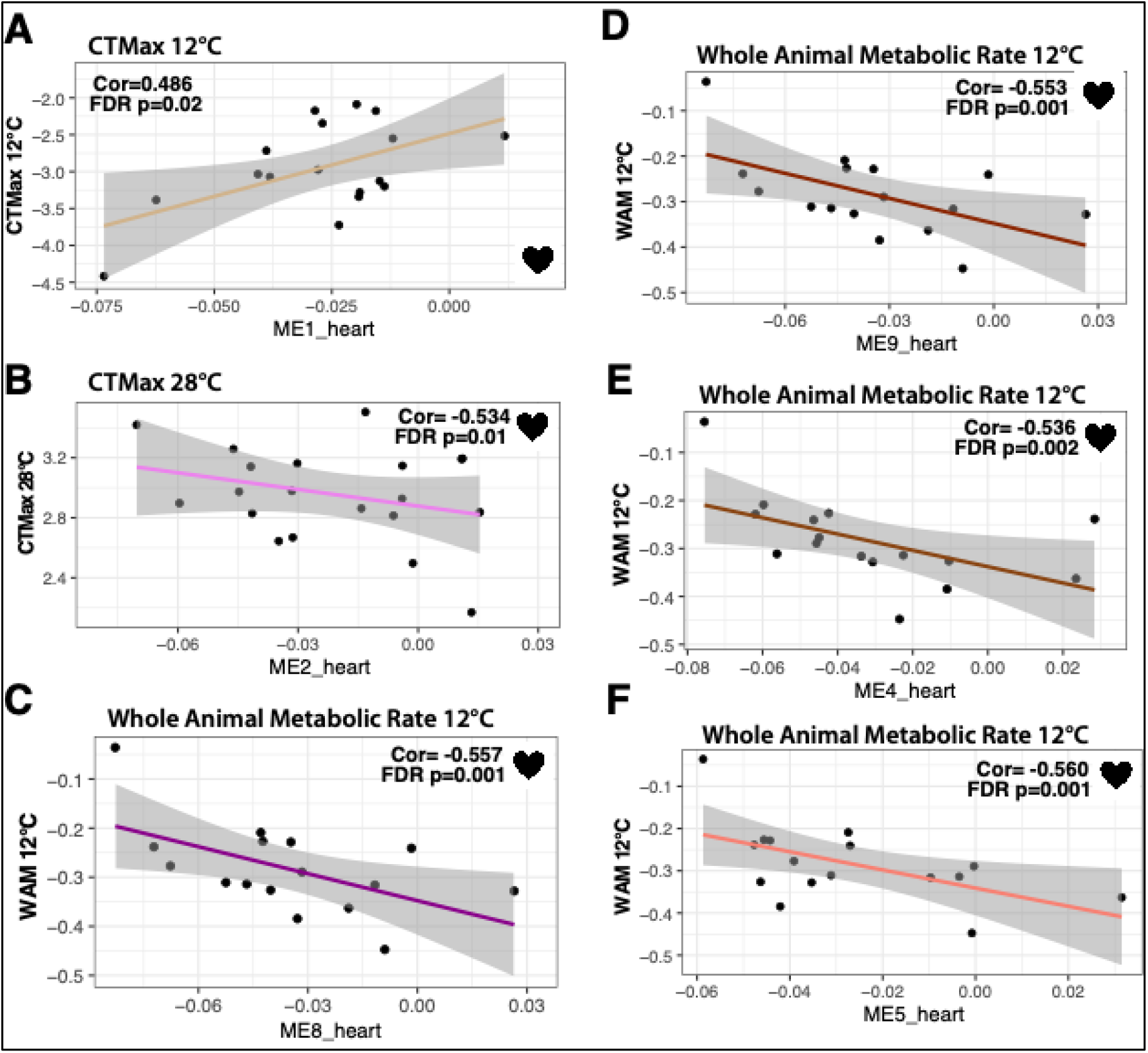
Significant whole animal trait-heart module correlations from weighted gene co-expression network analysis. Significant correlation of critical thermal maximum at 12°C (N=17) with ME1_heart (**A**), critical thermal maximum at 28°C (N=19) with ME2_heart (**B**), whole animal metabolic rate at 12°C (N=16) with ME8_heart (**C**), ME9_heart (**D**) ME4_heart (**E**), and ME5_heart (**F**). Pearson correlation coefficients (Cor) and FDR p-values are displayed for each significant correlation.

For traits that were significantly correlated with more than one ME, a multiple correlation coefficient was calculated. For WAM at 12°C, the four significant MEs together had a multiple correlation coefficient of 82%, the three significant MEs for FA CaM at 12°C had a multiple correlation coefficient of 79.5%, and the two significant MEs for LKA CaM at 12°C had a multiple correlation coefficient of 75.5%. All modules correlated with FA CaM at 12°C and CT_max_ at 12°C had positive correlation coefficients while all other significant trait *versus* ME correlations in hearts had negative correlation coefficients.

#### Brain WGCNA

Brain mRNAs had 42 total co-expressed modules with 142 to 393 mRNAs per module (Table 4). There were 6 significant ME-trait correlations (FDR <0.05) that included 4 unique modules and 4 temperature specific traits (Table 5, Fig. 5): at 12°C, body mass and at 28°C, CT_max_, WAM, and body mass. The trait with the most significant correlations was CT_max_ at 28°C (3 significant ME’s), and two of these were also significant with body mass at 28°C (ME3_brain) or WAM at 28°C (ME4_brain). On average, the correlation coefficient for a brain ME was 62% with a minimum of 56.8% (ME3_brain with CT_max_ 28°C) and a maximum of 70.2% (ME4_brain with WAM 28°C). For CT_max_ at 28°C, which was correlated with three MEs, the multiple correlation coefficient was 71.7%. All correlations between traits and brain MEs were negative except for body mass at 28°C.

**Figure 5:**
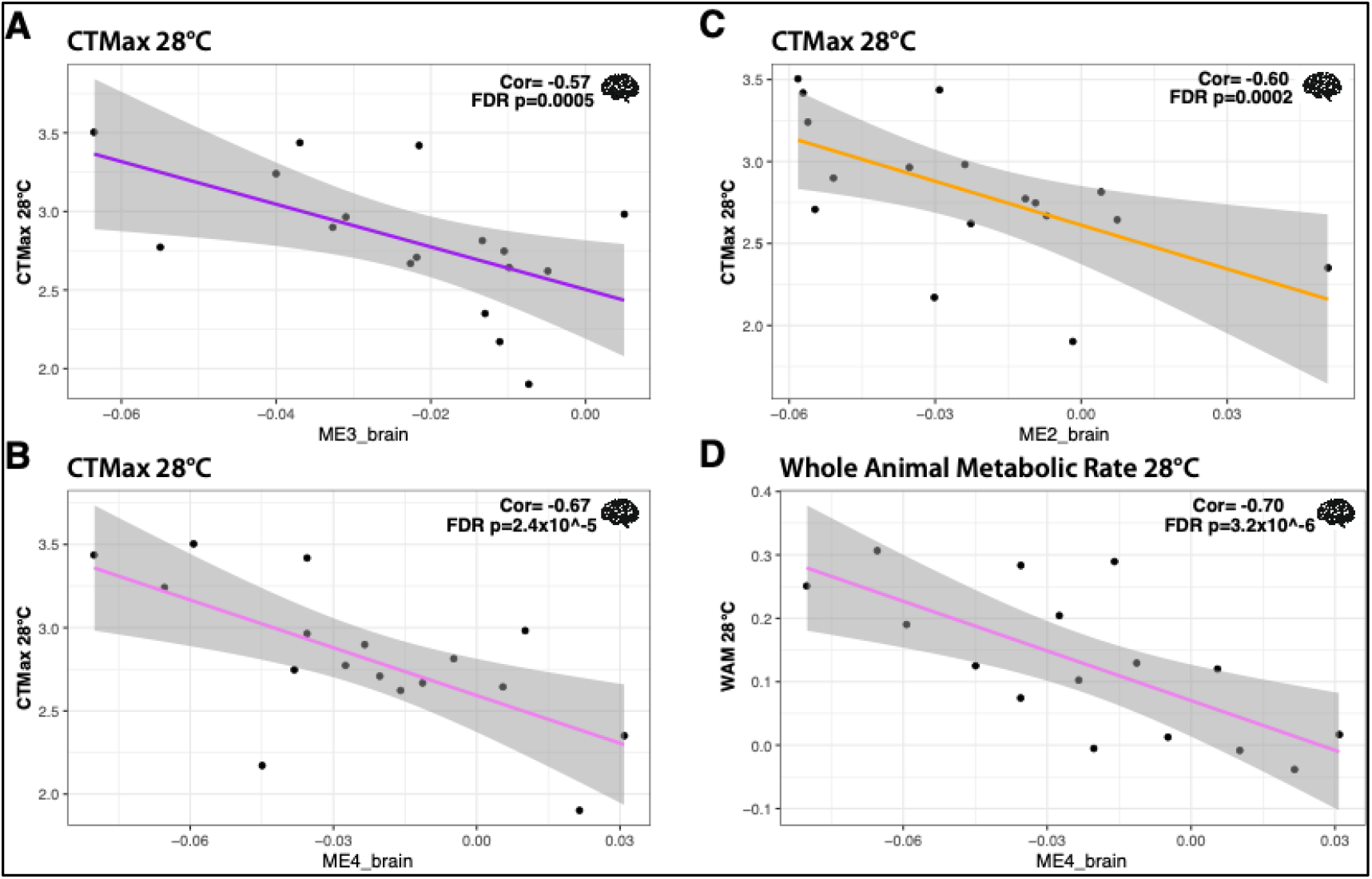
Significant whole animal trait-brain module correlations from weighted gene co-expression network analysis. Significant correlation of critical thermal maximum at 28°C (N=17) with ME3_brain (**A**), ME4_brain (**B**), and ME2_brain (**C**). Significant correlation of whole animal metabolic rate at 28°C (N=16) with ME4_brain (**D**). Pearson correlation coefficients (Cor) and FDR p-values are displayed for each significant correlation.

### KEGG and GO Enrichment

#### Critical thermal maximum enriched terms

MEs were tested for KEGG and GO term enrichment using the complete set of tissue specific mRNAs as the gene universal or reference set (10,535 for heart, 10,932 for brain; Tables S3 and S4). For CT_max_, all 5 MEs were significantly enriched for KEGG pathways (ME1_heart, ME2_heart, ME2_brain, ME3_brain, and ME4_brain) and included MAPK signaling, mTOR signaling, glyoxylate and dicarboxylate metabolism, insulin signaling pathway, glutathione metabolism, metabolic pathways, carbon metabolism, and tryptophan metabolism. Enriched GO terms included regulation of organ growth and cellular stress response in heart and AMP-activated protein kinase (AMPK) activity in brain. Notably, MAPK has been linked to adaptive cold tolerance (38, 39) and lipid metabolism (40). Additionally, mTOR is involved with energy homeostasis, has been linked to growth and longevity, and may be sensitive to temperature variation (41–43). Finally, AMPK induces cellular ATP production in mammals and is important for thermal stress response in ectotherms (44–46). Furthermore, AMPK phosphorylation in Coho salmon and rainbow trout hearts has been correlated with exposure above optimum temperatures (47), suggesting a role of AMPK in fish thermal response. ME for CT_max_ at 28°C contained substantial overlap in enriched KEGG pathways related to metabolism with 5 out of 8 terms enriched in both heart and brain modules. So, although there were different mRNAs in heart and brain modules correlated with CT_max_ at 28°C, the KEGG terms related to metabolism were shared (5 out of 8), suggesting that different mRNAs in heart and brain belonged to similar pathways impacting CT_max_ at 28°C.

#### Metabolic rate enriched terms

Modules significantly correlated with WAM were enriched for KEGG pathways including oxidative phosphorylation (heart only), glutathione metabolism (brain only), and metabolic pathways (both heart and brain). In addition, ME3_heart was correlated with FA CaM at 12°C and enriched for KEGG terms including metabolic pathways, forkhead protein (FoxO) signaling, and metabolism of NADH derivatives (nicotinate and nicotinamide). Notably, FoxO proteins, especially FoxO1, are involved in energy homeostasis and may aid in the switch from carbohydrate (glycolytic) to fatty acid metabolites (48). One module, ME5_heart, was significantly correlated with WAM at 12°C and FA CaM at 12°C and contained several KEGG pathways directly related to fatty acid metabolism as well as known transcription factors like PPAR that impact metabolic homeostasis by controlling expression of many metabolism related genes (49). Previously, partial correlation coefficients between FA CaM at 12°C and WAM at 12°C were negatively correlated (26), and similarly ME_5 had opposite correlation coefficients for these two traits (Table 3, Fig. 3, 4). This correlation between traits and their correlations to MEs also occur for CT_max_ and WAM at 28°C (26) and ME4_brain (enriched for glutathione metabolism and metabolic pathways). This emphasizes the biological relevance of the MEs in that the same MEs are associated with traits that have significant partial correlations.

## Discussion

### The role of mRNA expression in acclimation and evolution

For mRNA expression in both tissues, we found significant interactions between acclimation and population effects: the expression of several hundred mRNAs differed between acclimation temperatures, but these were not shared among all three populations (Fig. 1); also, the thermal effluent site (TE) had adaptive patterns different from the two reference sites that alter mRNA expression, but these differed between acclimation temperatures (Fig, S3). Previously (26), in these same individuals, acclimation response to 12°C and 28°C affected six physiological traits (WAM, CT_max_, and the four substrate specific CaM). For CT_max_, there was an expected enhancement: higher CT_max_ in individuals experiencing warmer environments. For metabolic rates (WAM and CaM), acclimation to 12°C and 28°C mitigated the effect of temperature (26). Specifically, without physiological acclimation there is an expected ∼3-fold increase in metabolic rates with the 16°C increase in acclimation and assay temperature (*i.e.,* with a doubling for every 10°C) (50). Yet, WAM had only ∼1.2-fold increase (34) from 12°C to 28°C, and CaM had no significant increase between temperatures (26). Presented here, across all three populations, acclimation produced significant differential mRNA expression (FDR <0.05) in hearts (366 mRNAs) and brains (528 mRNAs, (Table S1)). These mRNA expression changes associated with acclimation responses are similar to prior studies among ectotherms where transcriptomic response to temperature acclimation enhances thermal performance (34, 35, 39, 51). For example, in three-spine stickleback and other fishes, metabolic enzyme expression and mitochondrial volume density increase in response to cold acclimation can compensate for reduced enzyme catalytic rate with decreased temperature (34, 52, 53). Similarly, in eastern oysters (*Crassostrea virginica*), among a suite of environmental factors (temperature, pH, salinity, dissolved oxygen, etc.) temperature was the most important transcriptomic variation predictor with thermal stress increasing oxidative phosphorylation transcript expression (54). Even Antarctic fish, which are adapted to extreme cold, show plasticity in metabolic transcripts with temperature acclimation that impacts whole animal performance (53, 55).

While quantitative gene expression changes are common with acclimation (affecting both mRNA and proteins (39, 56)), what was surprising was that mRNA acclimation responses were different among populations—for heart mRNAs, no significant acclimation responses were shared among all three populations, and for brains only 1 mRNA was shared among the three populations. Further, 88-98% of significant acclimation responsive mRNAs are unique to each population. In contrast, the six physiological traits’ acclimation responses were not different among populations. All populations were subjected to a common environment for a long time (∼ 1 year or nearly 30-50% of a fish’s expected life span) with acclimation to the 12°C and 28C. Thus, the difference in acclimation mRNA response among populations was not due to short-term physiological effects and may be due to genetic polymorphisms driving acclimation responses but could also arise by irreversible developmental effects or trans-generational effects. Regardless of the genetic mechanisms responsible for the divergent mRNA acclimation responses among populations, these data suggest that multiple different mRNA expression patterns drive acclimation responses. This conclusion is similar to CaM measurements in Maine and Georgia populations, where the mRNAs that explain substrate specific metabolism varied among groups of individuals (6). The observations that plasticity in the six physiological traits between temperatures are similar among populations, yet mRNA acclimation responses differ among population, suggest that multiple redundant molecular mechanisms drive temperature compensation.

There is a single difference among populations for the six physiological traits: endogenous CaM at 28°C. Yet, populations had significant differences in mRNA expression specific for each temperature, and none of the population significant mRNAs were shared at 12°C or 28°C (Fig. S3, Table S2). One pattern, where the anthropogenically warmed TE population was significantly different from both northern and southern reference populations (not heated by thermal effluent from nuclear power plant), is indicative of adaptation (17, 32, 36). What differs here is that we examined mRNA expression that can be affected by DNA polymorphisms and also influenced by environment (i.e., GxE). Thus, the difference among populations in mRNA expression are dependent on the thermal environment, and if adaptive, suggest that the different genetic polymorphisms are responsible for adaptive divergence at different temperatures. This conclusion is similar to comparison within and among species: adaptive divergence in mRNA expression is dependent on the thermal environment experienced by individuals (17). For the TE population, Dayan *et al.,* conclude that there was adaptive divergence based on evolutionarily significant DNA polymorphisms (32). We would extend this to suggest that populations have evolved different mRNA expression patterns that are dependent on the thermal environment but that, nevertheless, produce similar physiological phenotypes.

### Biological relevance of co-expressed mRNAs

WGCNA identified co-expressed mRNA modules, MEs, highly correlated with WAM, CT_max_, FA CaM, LKA CaM, and body and heart mass, depending on the acclimation temperature (Fig, 3, 4, and 5). The average ME-trait correlation was 0.55 for heart and 0.62 for brains (Table 3 and 5). These MEs, containing 90-554 mRNAs each, contained few (0-10) mRNAs with significant expression differences among populations. WGCNA has been previously used to identify mRNA expression networks important for various pathologies including cardiovascular disease (57, 58), cancers (59–63), and diabetes (58, 64), among others. In non-human organisms, WGCNA has been used to characterize response to the environment, including heat stress in turbot (65), carotenoid metabolism in apricot fruit (66), and disease response in corals (67). Although few studies, to our knowledge, have validated correlations using jack-knife subsampling to ensure that the correlations were consistent among most individuals and not driven by a few outliers, these studies similarly identified potentially meaningful correlations between traits and co-expressed mRNAs. Importantly, in this study, the correlation patterns between MEs and physiological traits are similar to the correlations among physiological traits. For example at 12°C, FA CaM and WAM were negatively correlated (26) and similarly ME5_heart was significantly correlated with opposite signs with these two traits (*i.e.,* positively correlated with FA CaM but negatively correlated with WAM, Table 3, Fig. 3, 4). Additionally, MEs correlated to WAM and CaM were enriched in KEGG metabolic pathways and GO terms related to metabolism. These data indicate that modules represent independent, biologically important mRNA networks.

The biological importance of co-expressed mRNA networks is also supported by their relation to metabolic processes. Eleven of the 13 significant heart or brain MEs were significantly enriched for at least one KEGG term, and 6 were significantly enriched for at least one GO term. KEGG terms mapped to biologically relevant KEGG pathways including metabolic pathways, mechanistic target of rapamycin (mTOR) signaling, mitogen activated protein kinase (MAPK) signaling, insulin signaling, and metabolism and biosynthesis of various macromolecules including glycogen, NADH precursors, amino sugars, and fatty acids (Table S3, S4). Importantly, 8 out of 11 modules with significantly enriched KEGG terms mapped to at least one metabolism related KEGG pathway. Yet, we also found several enriched KEGG pathways and GO terms that were uniquely enriched in only one or few modules and seemingly unrelated to the correlated trait(s) (*e.g.,* cellular senescence). This could indicate a limited understanding of the complexity and interconnectedness among biological pathways and how different pathways affect a diversity of traits, mRNAs that are minimally annotated and missing relevant pathway involvement, or that mRNA expression impacts biological processes, that indirectly impact the traits we have measured. The concept that there is a limited understanding of the interactions among pathways is supported by mitochondrial respiration studies, specifically concerning the oxidative phosphorylation pathway (OxPhos) (68). When examining the selectively important nuclear genes effecting OxPhos, none of the genes were among the 97 proteins in the OxPhos pathway; instead, they were in diverse pathways, some of which made sense (*e.g,* ADP transport- where ADP is a substrate for OxPhos) (68). Thus, the few MEs associated with unexpected pathways may indicate a complexity in physiological traits where many pathways and the genes in these pathways affect trait variation.

Previously, data from our laboratory demonstrated that natural variation in substrate specific cardiac metabolism in *F. heteroclitus* could be explained by cardiac mRNA expression using microarray data (6). Similar to the WGCNA approach presented here, the first principal component of mRNA expression from different metabolic pathways (oxidative phosphorylation, TCA cycle, glycolysis) explained substrate specific CaM among individuals with different pathways of mRNAs explaining substrate specific metabolisms in different individuals. Here, we found that mRNA expression explained a similar proportion of substrate specific CaM as previously reported (∼80%) using three MEs to explain a single trait (FA CaM at 12°C).

Few, if any, studies have examined the correlation of co-expressed mRNA with CT_max_ (although see (69)). Our analyses found that 341 heart mRNAs in two co-expressed modules and 682 brain mRNAs in three co-expressed modules were associated with CT_max_ at 12°C or 28°C, with different MEs at each temperature. Futhermore, heart and brain significant ME for CT_max_ at 28°C share enriched KEGG pathways, yet do not share any mRNAs, suggesting that different mRNAs affect a common set of pathways that impact CTmax. These data suggest that CTmax is polygenic and relies on different mRNAs in different tissues at different temperatures. There is prior evidence suggesting that CT_max_ is polygenic: a GBS study covering ∼ 0.1% of the genome found up to 47 single nucleotide polymorphisms (SNPs) that explained 43.4% of variation in CT_max_ among *F. heteroclitus* individuals collected from Georgia, New Jersey, and New Hampshire, USA (70). Here, a greater proportion of CT_max_ was explained with mRNA expression, up to 71.7% with 3 brain MEs. This increase in explained CT_max_ variance by mRNA expression is likely due to the combined heritable and physiologically inducible nature of mRNA expression. Few (0-10) of the mRNAs in MEs were differentially expressed between 12°C and 28°C, and thus MEs that explained CT_max_ variation within each of acclimation temperatures are not due to acclimation effects on mRNA expression. Instead, the CT_max_ variation within each acclimation temperature appears to be due to individual variation in mRNA expression, which may be explained by nucleotide variation driving differential expression.

Whole animal metabolism, WAM, is a fundamental physiological process that defines how animals live, niche space, evolutionary transition, and the human condition (71–75). There is significant literature investigating metabolic rate variation (*e.g.* (31, 76, 77)); however, the relationship between metabolic rate and mRNA expression remains poorly understood. This may be due to the complex nature of whole animal metabolism, which is a sum of tissue specific metabolic demands and a balance between growth, maintenance, and energy storage. Yet, we find 82% of 12°C WAM variation related to four heart MEs with 736 mRNAs and 50% of 28°C WAM variation related to one brain ME with 142 mRNAs. These data indicate that a large proportion of WAM can be explained by mRNAs within a common pathway impacting cardiac metabolic processes and thus provides insight into the physiological relationship between cardiorespiratory performance and overall metabolism (78–81).

These WGCNA analyses suggest that many mRNAs in several biochemical pathways define the physiological state among individuals. Yet, the careful reader will note two substantial complexities: 1) heart MEs explain the variation in many physiological traits at 12°C but few at 28°C, and brain MEs explain the variation in many physiological traits at 28°C but few at 12°C and 2) mRNAs within MEs are enriched for many diverse and unexpected pathways as discussed above.

The difference between tissue specific MEs and their association with physiological traits is related to CaM, WAM, and CT_max_ having higher inter-individual variation at 12°C than 28°C, and similarly there is greater mRNA expression variation at 12°C than at 28°C. Thus, the more frequent explanation of physiological traits by 12°C mRNA expression may simply result from greater statistical power due to the greater variance in both physiological traits and mRNA. Yet, in brains, mRNAs explain WAM and CT_max_ at 28°C. While we can only speculate, these data suggest that at the higher temperature brain mRNA expression is more important than cardiac mRNA expression. There is evidence that acclimatory response to temperature in brain is greater than in hearts (more acclimatory mRNAs and greater decreased mitochondrial function in brain when compared to heart tissue (82)). This is similar to our data: brains at 28°C have more acclimation responsive mRNAs than hearts, and more population divergence than brains at 12°C or hearts at either temperature. Together these data suggest that the variation in WAM and CTmax are more dependent on brain specific expression at 28°C.

Many of MEs are associated with KEGG pathways that impinge on metabolic processes. Physiological processes are within 9 heart MEs and 4 brain MEs (Tables 2 and 4), and these ME each contain 90-554 mRNAs. Each of these MEs is significantly enriched for multiple KEGG and GO pathways (Tables S3 and S4), including pathways not typically thought to be directly involved in metabolism or thermal tolerance. Similar to nuclear genes that impact *Fundulus* mitochondrial respiration (68), these data suggest that many metabolically distant genes affect physiological variation. This is important because too often publications have “just so stories” (a la (83, 84)) that only focus on a few preconceived genes to explain functional physiological variation (85). While it is understandable to highlight a prior expectation, doing so limits our understanding of how genotypes effect phenotypes.

In summary, the data provided here build on previous findings that inter-individual mRNA expression variation is biologically important by identifying that 1) mRNAs important for acclimation are population specific, 2) divergence among geographically close populations does not include acclimation responsive mRNAs, and 3) mRNA expression associated with physiological traits is tissue specific and dependent on the thermal environment. We highlight that biologically important mRNA networks are related to 48-82% of variation in whole animal metabolism thermal tolerance, or substrate specific cardiac metabolism and are different at different thermal environments. This suggests that mRNA variation among individuals within and among populations is important for explaining complex trait variation and, surprisingly, that while similar pathways can be important at different temperatures, the tissues where they are expressed differ: heart mRNA expression explains variation in more traits at 12°C, and brain mRNA explains variation in more traits at 28°C.

## Methods

### Animal care and use

Fish were collected in live traps in September 2018 at three sites in central New Jersey, USA near the Oyster Creek Nuclear generating station (OCNGS). Sites included one north reference (N.Ref; 39°52’28.000 N, 74°08’19.000 W), one south reference (S.Ref; 39°47’04.000 N, 74°11’07.000 W), and a central site located within the thermal effluent of the OCNGS (TE; 39°48’33.000 N, 74°10’51.000 W). All fish were transferred live to the University of Miami, FL where they were kept in accordance with the University of Miami Institutional Animal Care and Use Committee (IACUC) guidelines.

Individuals from all three populations were common gardened to 20°C for three months (12:12 light dark cycle) and kept at 20°C in a common recirculating seawater system (15ppt) at 12 hours light:12 hours dark, then subjected to *pseudo-*winter for 6 weeks at 8°C (8:16 light dark cycle). Following the *pseudo-winter*, half of the fish from each population were acclimated to 12°C and the other half to 28°C (16:8 light dark cycle) for four weeks prior to determination of WAM and CT_max_. Following this acclimation, fish originally acclimated to 12°C were acclimated to 28°C and *vice versa* for at least four weeks, and WAM and CT_max_ were measured at the new acclimation temperature. After a minimum one-week recovery period post-CT_max_, fish were sacrificed, substrate specific CaM was measured at the second acclimation temperature, and mRNA was isolated. Thus, WAM and CT_max_ were measured in all individuals at 12°C and 28°C, but CaM and mRNA were sampled from half the individuals acclimated at 12°C and the other half at 28°C.

### Quantifying metabolic and thermal tolerance traits

Individuals acclimated to 12°C and 28°C for at least 4 weeks were measured for whole animal metabolic rate (overnight intermittent flow respirometry) with a minimum of 20 replicate metabolic rate measures per individual used to determine the standard metabolic rate (SMR) in mgO2 hr^-1^ (7). Critical thermal maximum (CT_max_) was measured in a 10-gallon tank that was slowly heated at a rate of 0.3°C min^-1^ as in (86) and was defined as the point when fish lost equilibrium in the water column for 5 consecutive seconds. Finally, substrate specific cardiac metabolic rate (CaM, substrates: 5mM glucose, fatty acids – 1 mM Palmitic acid conjugated to fatty-acid-free bovine serum albumin, lactate+ketones+ethanol – 5 mM lactate, 5 mM hydroxybutyrate, 5 mM ethyl acetoacetate, 0.1% ethanol, endogenous – substrate free Ringers media) was measured using micro-respirometry. Heart ventricles were dissected out and splaying in Ringers media before being transferred to a custom 1mL chamber system (10). Each heart was measured for glucose (GLU), then fatty acids (FA), followed by lactate+ketones+ethanol (LKA), and endogenous (END) cardiac metabolic rate. All substrates except GLU used non-reversible glycolytic enzyme inhibitors, so the order of substrates did not differ among hearts. Each heart was measured for 6 minutes per substrate with the last 3 minutes used to calculated oxygen consumption in pmolO_2_ sec^-1^. All respirometry measurements used Presens fiber optic oxygen sensors with flow through cells (WAM) or sensor spots (CaM). Both WAM and CaM were measured at both acclimation temperatures (12°C and 28°C) for the same individuals, and CaM was measured at a single acclimation temperature. For additional methods and analysis of physiological data see (26).

### mRNA library preparation and sequencing

Tissues for mRNA expression were stored in chaotropic buffer at the time of CaM measurement and captured gene expression variation due to long term temperature acclimation rather than heat shock or acute temperature response (*e.g.,* resulting from higher temperatures during CT_max_ measurements) because CT_max_ measurements were performed at least a week prior to tissue isolation. We extracted total RNA from homogenized heart and brain tissues using phenol-chloroform isoamyl alcohol isolation and treated RNA samples with DNAse to remove genomic DNA. For each sample we started with 50ng of RNA and captured the 3’ mRNA ends using an NVdT primer with a poly-A tail for first strand cDNA synthesis (Table S5). This primer contained a unique barcode for each sample (1–96), which allowed all samples in a single plate to be pooled for the remaining library preparation steps. Nick translation was used to make double stranded cDNA that was digested with an in-house purified Tn5 transposase (as in (87)) loaded with partial adapter sequences to generate fragments of double stranded cDNA ranging from ∼300-800bp (Table S6). Libraries were amplified for 17 PCR cycles using primers complimentary to the inserted partial adapter sequence and a plate level barcode to fully multiplex samples.

### mRNA data processing and analysis

A total of 219 libraries (110 individuals, 2 tissues per individual, 1 individual only heart was collected) were pooled and sequenced on 2 lanes of Illumina HiSeq4000 (dual end 150bp reads) at the Genewiz LCC facility, South Plainfield NJ, USA. Raw reads were trimmed with BBDuk (from BBMap v. 38.87) to remove adapter sequences, aligned with STAR (v. 2.7.5) to the *Fundulus heteroclitus* genome, and counted with Featurecounts (v. 1.4.6-p5, parameters: -T 4 -s 2 -t gene -g gene_id).

The raw counts table was imported into R Studio (v. 1.4.1106) and all counts were normalized for library size using the median ratio method (88) with the “estimateSizeFactors” function in DESeq2 (33). Samples with a minimum of 1.5 million reads for hearts or 1 million reads for brains were retained and filtered to keep only mRNAs with at least 30 counts in 10% of individuals. Principal component analysis (PCA) using “plotPCA” function from the DESeq2 package was used to examine variation among all samples. Among all samples, the 500 most variable mRNAs were used for PCA. In this analysis, 12 hearts and 13 brains were removed as outliers because they differed in expression from other same-tissue samples (*i.e.,* some hearts had “brain-like” expression patterns and *vice versa,* Fig. S1). This reduced variation within a tissue and reduced sample size; however, the individuals removed were not from a single acclimation temperature or population so likely had little overall impact on further analyses. In a separate tissue specific principal component analysis, clustering of samples by biological effects including sex, habitat temperature, population, acclimation temperature, and date of tissue collection (possible batch effect) was examined. Batch effects did not split individuals along any of the principal components examined (PC1-PC4) for heart or brain. In this separate analysis, heart PC1 accounted for 18%, heart PC2 for 7%, brain PC1 for 11%, and brain PC2 for 7% of the variation among individuals. No biologically relevant clustering was detected among the first 4 principal components for either tissue (chi-squared test p>0.05, Fig. S2A, B). To determine the degree of variation in mRNA expression among groups within a tissue, the coefficient of variation (CV, standard deviation/mean) for each expressed mRNA was calculated and the average CV compared among groups.

### Differential expression analysis

DESeq2 (33) package in R was used for differential expression analysis separately for heart and brain. To identify differentially expressed mRNAs between acclimation temperatures within populations, the DESeq model used was: ∼Population + Acclimation_Temperature + Population*Acclimation_Temperature. Additionally, due to significant interaction between population and acclimation temperature, a separate analysis was used to find differentially expressed mRNAs among populations within an acclimation temperature; individuals measured for CaM only at 12°C or 28°C were used with DESeq model: ∼Population. Multiple test correction across all comparisons made within a model used the Benjamin Hochberg false discovery rate with a significance threshold of 0.05.

### Weighted gene co-expression network analysis

To identify sets of co-expressed mRNAs, weighted gene co-expression network analysis (WGCNA, (37)) was completed for heart and brain separately. Network calculation, used to group mRNAs into co-expressed modules, for heart and brain, used soft thresholding to generate a scale free network with high similarity (soft thresholding power set to 5 in heart, 4 in brain) before calculating the topological overlap measure (TOM) and using dynamicTree with minimum module size of 30 and threshold for module merging of 0.75. The first principal component of each independent module (below the threshold for module merging), known as the module eigengene (ME), was then correlated with temperature specific quantitative traits using signed Pearson’s correlation. Multiple test correction across all correlations made for a single trait used the Benjamin Hochberg false discovery rate with a significance threshold of 0.05. To remove significant correlations potentially driven by outliers, a jack-knife approach was used to subsample 90% of individuals and repeat the signed Pearson correlation analysis 100 times. Correlations that were significant in >70 out of 100 subsamples in the same direction were robust to outliers and reported as significant. A multivariate correlation coefficient was calculated for traits significantly correlated with more than one module by correlating the fitted values from a linear model with formula: trait∼ME1+ME2..ME# with trait data. This multivariate correlation coefficient represents the ability of the MEs together to accurately predict the trait. For significant modules, the mRNAs with the highest module membership (MM, correlation between mRNA expression and module eigengene) and the highest gene significance for a given trait within a module (GS, correlation between mRNA expression and a given quantitative trait) were also identified.

### Gene Ontology and Kyoto Encyclopedia of Genes and Genomes Enrichment

To identify biologically important networks within WGCNA modules that were significantly correlated with at least 1 trait, we used Kyoto Encyclopedia of Genes and Genomes (KEGG) pathway and gene ontology (GO) enrichment analyses. First, the genome was mapped to KEGG and GO terms using eggNOG mapper (89) with default parameters. The KEGG and GO terms were then matched to the set of expressed mRNAs for heart and brain. The list of mRNAs in each module was compared to the set of expressed mRNAs (set as the reference or gene universe) in each tissue for enrichment analysis in R using the clusterProfiler package “enricher” function for KEGG terms (90). To map enriched KEGG terms to KEGG pathways, the KEGG Mapper online tool was used with annotations from the closest relative, zebrafish (*Danio rerio*) (91, 92). Cytoscape BiNGO was used for GO enrichment using the set of expressed mRNAs as the reference to examined enrichment of biological process, molecular function, and cellular component GO terms (93). Significant KEGG and GO terms are reported with FDR p-value threshold of 0.05.

## Data Availability

All sequence data is available in NCBI SRA: https://dataview.ncbi.nlm.nih.gov/object/PRJNA796010?reviewer=f1u5dbo46558lklv8m2cknap87 (public DOI available upon publication). All physiological data is available in Dryad: https://doi.org/10.5061/dryad.0gb5mkm0w. Code for processing of raw sequence files and all data analysis and visualization conducted in R is available in Github: https://github.com/mxd1288/OCNJ_F18_RNA.git.

## Author Contributions

Data collection, analysis, and visualization MKD. Original manuscript draft MKD. Manuscript editing MFO and DLC. Study design MKD, MFO, and DLC. This research was supported by funding from National Science Foundation award nos. IOS 1556396 and IOS 1754437 to MFO and DLC.

## Acknowledgements

Thank you to Amanda DeLiberto and Samantha Sierra-Martinez for assistance in isolation of the Tn5 enzyme used for RNA library preparation and to Dr. Benjamin Young for support in RNA data analysis.

## Supplemental Figures

**Figure S1:**
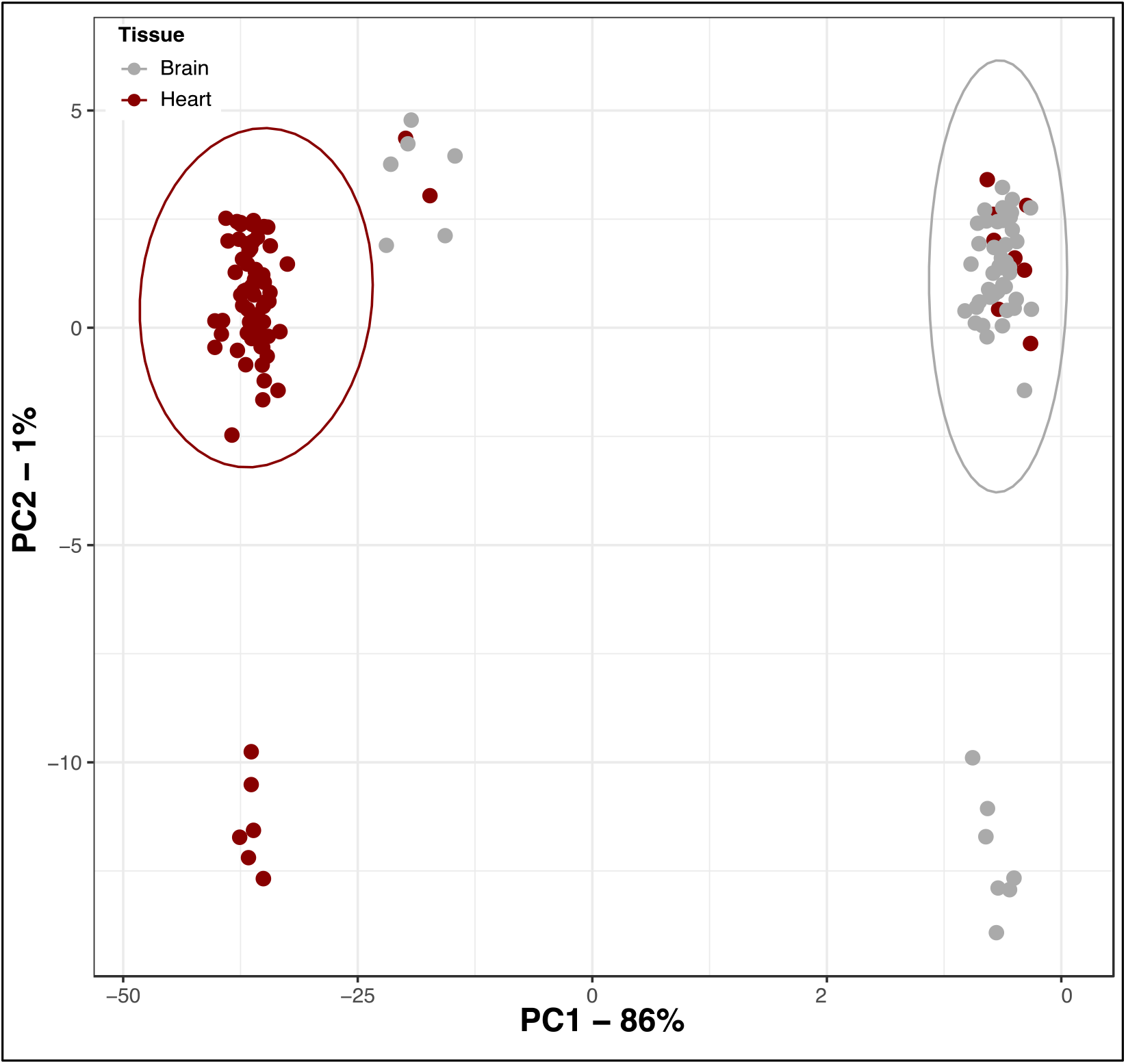
Principal component analysis of all samples. Principal component 1 split heart and brain tissue and explained 86% of variance. Individuals who did not clearly group with the appropriate tissue were removed as outliers.

**Figure S2:**
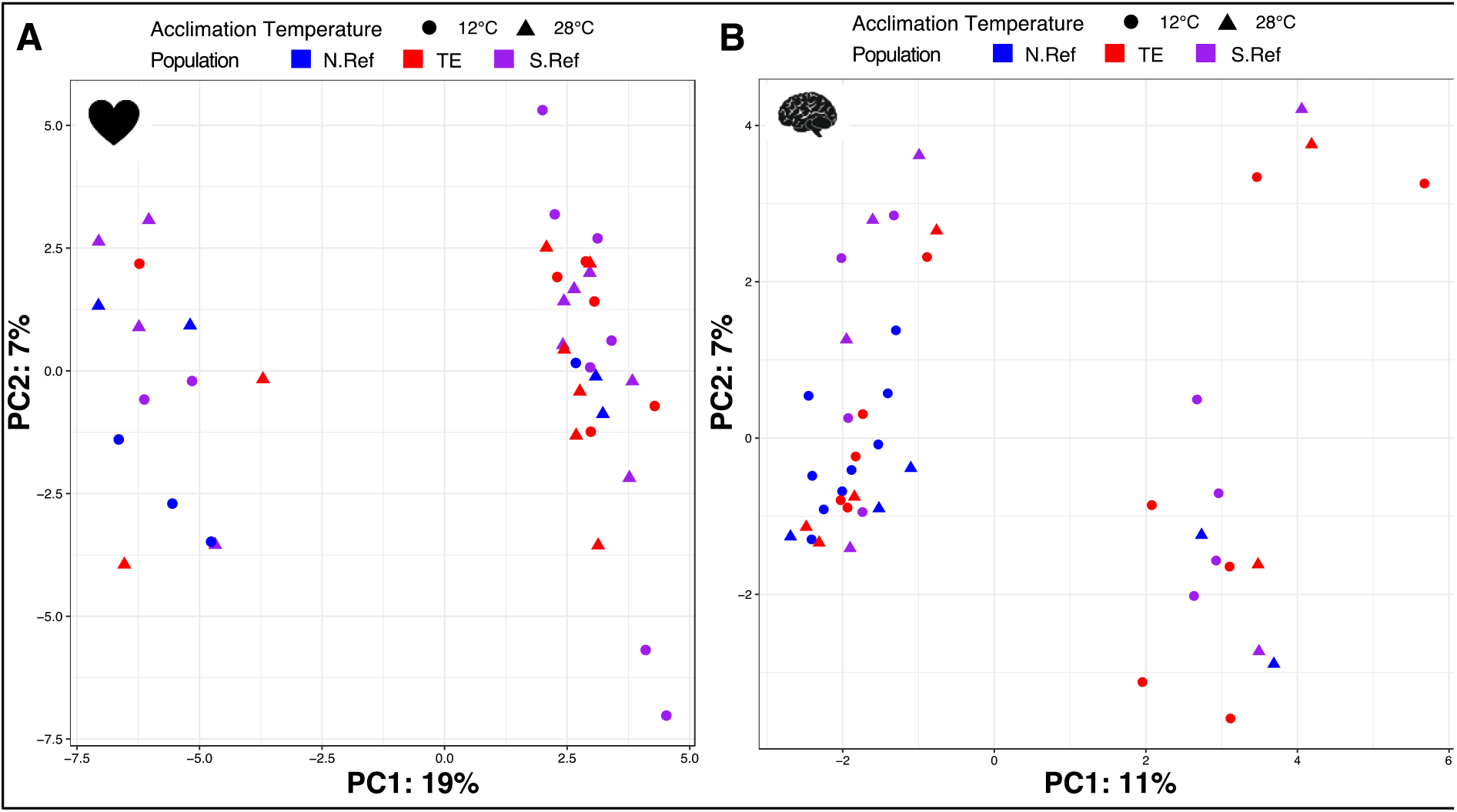
Tissue specific principal component analysis. Heart (N=41, A) first two principal components explain 19% and 7% of variance. Brain (N=45, B) first two principal components explain 11% and 7% of variance. Triangles are 28°C acclimated individuals, circles are 12°C. Individuals from the north reference (N.Ref) are blue, south reference (S.Ref) are purple, and thermal effluent population (TE) are red.

**Figure S3:**
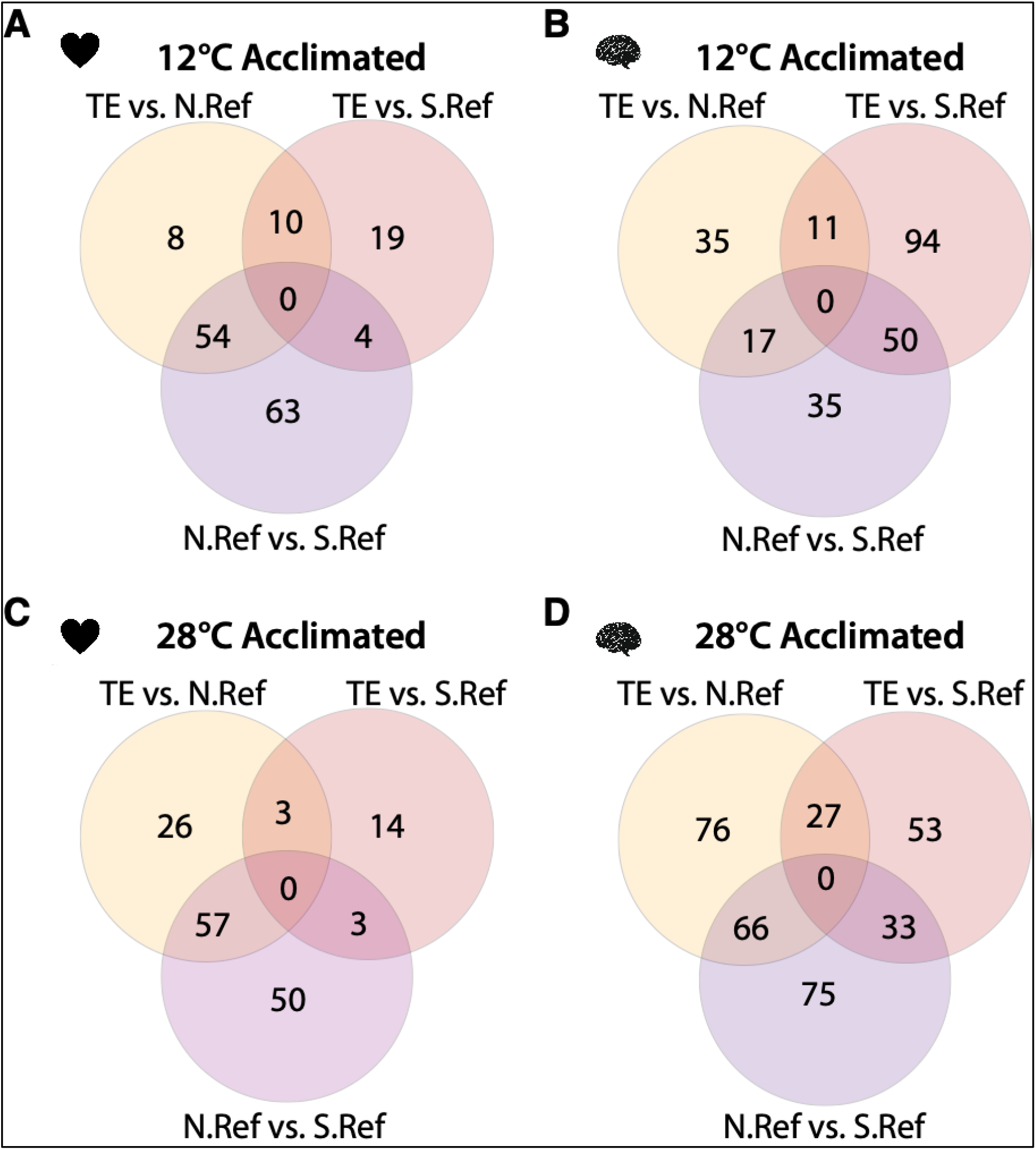
Differentially expressed mRNAs among populations within tissue and acclimation temperature. **A)** Heart at 12°C, **B)** brain at 12°C, **C)** heart at 28°C, D) brain at 28°C. Populations are north reference (N.Ref), south reference (S.Ref), and thermal effluent (TE).

## References

1. Albert FW, Kruglyak L. The role of regulatory variation in complex traits and disease. Nat Rev Genet. 2015;16(4):197–212.

2. Morgante F, Huang W, Sorensen P, Maltecca C, Mackay TFC. Leveraging Multiple Layers of Data To Predict Drosophila Complex Traits. G3 (Bethesda). 2020;10(12):4599–613.

3. Gao AWW, Sterken MG, de Bos JU, van Creij J, Kamble R, Snoek BL, et al. Natural genetic variation in C. elegans identified genomic loci controlling metabolite levels. Genome Res. 2018;28(9):1296–308.

4. Marchadier E, Hanemian M, Tisne S, Bach L, Bazakos C, Gilbault E, et al. The complex genetic architecture of shoot growth natural variation in Arabidopsis thaliana. Plos Genet. 2019;15(4).

5. Crawford DL, Oleksiak MF. The biological importance of measuring individual variation. J Exp Biol. 2007;210(9):1613–21.

6. Oleksiak MF, Roach JL, Crawford DL. Natural variation in cardiac metabolism and gene expression in Fundulus heteroclitus. Nature Genetics. 2005;37(1):67–72.

7. Drown MK, DeLiberto AN, Crawford DL, Oleksiak MF. An Innovative Setup for High-Throughput Respirometry of Small Aquatic Animals. Frontiers in Marine Science. 2020;7.

8. Christensen EAF, Andersen LEJ, Bergsson H, Steffensen JF, Killen SS. Shuttle-box systems for studying preferred environmental ranges by aquatic animals. Conserv Physiol. 2021;9(1):coab028.

9. Yang WN, Feng H, Zhang XH, Zhang J, Doonan JH, Batchelor WD, et al. Crop Phenomics and High-Throughput Phenotyping: Past Decades, Current Challenges, and Future Perspectives. Mol Plant. 2020;13(2):187–214.

10. DeLiberto AN, Drown MK, Oleksiak MF, Crawford DL. Measuring complex phenotypes: A flexible high-throughput design for micro-respirometry. bioRxiv. 2020.

11. Barghi N, Tobler R, Nolte V, Jakšić AM, Mallard F, Otte KA, et al. Genetic redundancy fuels polygenic adaptation in Drosophila. PLOS Biology. 2019;17(2):e3000128.

12. Laruson AJ, Yeaman S, Lotterhos KE. The Importance of Genetic Redundancy in Evolution. Trends in ecology & evolution. 2020.

13. Yeaman S. Local Adaptation by Alleles of Small Effect. Am Nat. 2015;186:S74–S89.

14. Gibson G, Weir B. The quantitative genetics of transcription. Trends Genet. 2005;21(11):616–23.

15. Kliebenstein DJ. A role for gene duplication and natural variation of gene expression in the evolution of metabolism. PLoS One. 2008;3(3):e1838.

16. McCairns RJ, Bernatchez L. Adaptive divergence between freshwater and marine sticklebacks: insights into the role of phenotypic plasticity from an integrated analysis of candidate gene expression. Evolution. 2010;64(4):1029–47.

17. Dayan DI, Crawford DL, Oleksiak MF. Phenotypic plasticity in gene expression contributes to divergence of locally adapted populations of Fundulus heteroclitus. Molecular Ecology. 2015;24(13):3345–59.

18. Levine MT, Eckert ML, Begun DJ. Whole-genome expression plasticity across tropical and temperate Drosophila melanogaster populations from Eastern Australia. Mol Biol Evol. 2011;28(1):249–56.

19. Kvist J, Wheat CW, Kallioniemi E, Saastamoinen M, Hanski I, Frilander MJ. Temperature treatments during larval development reveal extensive heritable and plastic variation in gene expression and life history traits. Mol Ecol. 2013;22(3):602–19.

20. Fay JC, McCullough HL, Sniegowski PD, Eisen MB. Population genetic variation in gene expression is associated with phenotypic variation in Saccharomyces cerevisiae. Genome Biol. 2004;5(4):R26.

21. Calderwood A, Lloyd A, Hepworth J, Tudor EH, Jones DM, Woodhouse S, et al. Total FLC transcript dynamics from divergent paralogue expression explains flowering diversity in Brassica napus. New Phytol. 2021;229(6):3534–48.

22. Yechoor VK, Patti ME, Saccone R, Kahn CR. Coordinated patterns of gene expression for substrate and energy metabolism in skeletal muscle of diabetic mice. Proc Natl Acad Sci U S A. 2002;99(16):10587–92.

23. Middleton FA, Mirnics K, Pierri JN, Lewis DA, Levitt P. Gene expression profiling reveals alterations of specific metabolic pathways in schizophrenia. J Neurosci. 2002;22(7):2718–29.

24. Rockman MV, Kruglyak L. Genetics of global gene expression. Nat Rev Genet. 2006;7(11):862–72.

25. Hill MS, Vande Zande P, Wittkopp PJ. Molecular and evolutionary processes generating variation in gene expression. Nat Rev Genet. 2021;22(4):203–15.

26. Drown MK, DeLiberto AN, Ehrlich MA, Crawford DL, Oleksiak MF. Interindividual plasticity in metabolic and thermal tolerance traits from populations subjected to recent anthropogenic heating. R Soc Open Sci. 2021;8(7):210440.

27. Morgan R, Finnoen MH, Jutfelt F. CTmax is repeatable and doesn’t reduce growth in zebrafish. Sci Rep. 2018;8(1):7099.

28. Ronning B, Jensen H, Moe B, Bech C. Basal metabolic rate: heritability and genetic correlations with morphological traits in the zebra finch. Journal of evolutionary biology. 2007;20(5):1815–22.

29. Mattila AL, Hanski I. Heritability of flight and resting metabolic rates in the Glanville fritillary butterfly. Journal of evolutionary biology. 2014;27(8):1733–43.

30. Nilsson JA, Akesson M, Nilsson JF. Heritability of resting metabolic rate in a wild population of blue tits. Journal of evolutionary biology. 2009;22(9):1867–74.

31. Reemeyer JE, Rees BB. Plasticity, repeatability and phenotypic correlations of aerobic metabolic traits in a small estuarine fish. J Exp Biol. 2020;223(Pt 14).

32. Dayan DI, Du X, Baris TZ, Wagner DN, Crawford DL, Oleksiak MF. Population genomics of rapid evolution in natural populations: polygenic selection in response to power station thermal effluents. BMC Evol Biol. 2019;19(1):61.

33. Love MI, Huber W, Anders S. Moderated estimation of fold change and dispersion for RNA-seq data with DESeq2. Genome Biol. 2014;15(12):550.

34. Orczewska JI, Hartleben G, O’Brien KM. The molecular basis of aerobic metabolic remodeling differs between oxidative muscle and liver of threespine sticklebacks in response to cold acclimation. Am J Physiol Regul Integr Comp Physiol. 2010;299(1):R352–64.

35. Podrabsky JE, Somero GN. Changes in gene expression associated with acclimation to constant temperatures and fluctuating daily temperatures in an annual killifish Austrofundulus limnaeus. J Exp Biol. 2004;207(Pt 13):2237–54.

36. Williams LM, Oleksiak MF. Signatures of selection in natural populations adapted to chronic pollution. BMC Evol Biol. 2008;8:282.

37. Langfelder P, Horvath S. WGCNA: an R package for weighted correlation network analysis. BMC Bioinformatics. 2008;9:559.

38. Chen ZZ, Cheng CHC, Zhang JF, Cao LX, Chen L, Zhou LH, et al. Transcrintomic and genomic evolution under constant cold in Antarctic notothenioid fish. P Natl Acad Sci USA. 2008;105(35):12944–9.

39. Wen X, Zhang XY, Hu YD, Xu JJ, Wang T, Yin SW. iTRAQ-based quantitative proteomic analysis of Takifugu fasciatus liver in response to low-temperature stress. J Proteomics. 2019;201:27–36.

40. Bost F, Aouadi M, Caron L, Binetruy B. The role of MAPKs in adipocyte differentiation and obesity. Biochimie. 2005;87(1):51–6.

41. Aramburu J, Ortells MC, Tejedor S, Buxade M, Lopez-Rodriguez C. Transcriptional regulation of the stress response by mTOR. Sci Signal. 2014;7(332).

42. Sengupta S, Peterson TR, Laplante M, Oh S, Sabatini DM. mTORC1 controls fasting-induced ketogenesis and its modulation by ageing. Nature. 2010;468(7327):1100–U502.

43. Reiling JH, Sabatini DM. Stress and mTORture signaling. Oncogene. 2006;25(48):6373–83.

44. Seebacher F, Little AG. Plasticity of Performance Curves Can Buffer Reaction Rates from Body Temperature Variation in Active Endotherms. Front Physiol. 2017;8:575.

45. Wittmann AC, Benrabaa SAM, Lopez-Ceron DA, Chang ES, Mykles DL. Effects of temperature on survival, moulting, and expression of neuropeptide and mTOR signalling genes in juvenile Dungeness crab (Metacarcinus magister). Journal of Experimental Biology. 2018;221(21).

46. Frederich M, O’Rourke MR, Furey NB, Jost JA. AMP-activated protein kinase (AMPK) in the rock crab, Cancer irroratus: an early indicator of temperature stress. Journal of Experimental Biology. 2009;212(5):722–30.

47. Anttila K, Casselman MT, Schulte PM, Farrell AP. Optimum Temperature in Juvenile Salmonids: Connecting Subcellular Indicators to Tissue Function and Whole-Organism Thermal Optimum. Physiol Biochem Zool. 2013;86(2):245–56.

48. Gross DN, van den Heuvel APJ, Birnbaum MJ. The role of FoxO in the regulation of metabolism. Oncogene. 2008;27(16):2320–36.

49. Han L, Shen WJ, Bittner S, Kraemer FB, Azhar S. PPARs: regulators of metabolism and as therapeutic targets in cardiovascular disease. Part I: PPAR-alpha. Future Cardiol. 2017;13(3):259–78.

50. Schulte PM, Healy TM, Fangue NA. Thermal performance curves, phenotypic plasticity, and the time scales of temperature exposure. Integr Comp Biol. 2011;51(5):691–702.

51. Zhou S, Campbell TG, Stone EA, Mackay TF, Anholt RR. Phenotypic plasticity of the Drosophila transcriptome. Plos Genet. 2012;8(3):e1002593.

52. Fangue NA, Richards JG, Schulte PM. Do mitochondrial properties explain intraspecific variation in thermal tolerance? J Exp Biol. 2009;212(Pt 4):514–22.

53. Lucassen M, Schmidt A, Eckerle LG, Portner H-O. Mitochondrial proliferation in the permanent vs. temporary cold: enzyme activities and mRNA levels in Antarctic and temperate zoarcid fish. American Journal of Physiology-Regulatory, Integrative and Comparative Physiology. 2003;285(6):R1410–R20.

54. Chapman RW, Mancia A, Beal M, Veloso A, Rathburn C, Blair A, et al. The transcriptomic responses of the eastern oyster, Crassostrea virginica, to environmental conditions. Molecular Ecology. 2011;20(7):1431–49.

55. Windisch HS, Frickenhaus S, John U, Knust R, Pörtner H-O, Lucassen M. Stress response or beneficial temperature acclimation: transcriptomic signatures in Antarctic fish (Pachycara brachycephalum). Molecular Ecology. 2014;23(14):3469–82.

56. Somero GN. Proteins and temperature. Annu Rev Physiol. 1995;57:43–68.

57. Tang Y, Ke ZP, Peng YG, Cai PT. Co-expression analysis reveals key gene modules and pathway of human coronary heart disease. J Cell Biochem. 2018;119(2):2102–9.

58. Liang W, Sun F, Zhao Y, Shan L, Lou H. Identification of Susceptibility Modules and Genes for Cardiovascular Disease in Diabetic Patients Using WGCNA Analysis. J Diabetes Res. 2020;2020:4178639.

59. Di Y, Chen D, Yu W, Yan L. Bladder cancer stage-associated hub genes revealed by WGCNA co-expression network analysis. Hereditas. 2019;156:7.

60. Zhai X, Xue Q, Liu Q, Guo Y, Chen Z. Colon cancer recurrenceassociated genes revealed by WGCNA coexpression network analysis. Mol Med Rep. 2017;16(5):6499–505.

61. Su R, Jin C, Zhou L, Cao Y, Kuang M, Li L, et al. Construction of a ceRNA network of hub genes affecting immune infiltration in ovarian cancer identified by WGCNA. BMC Cancer. 2021;21(1):970.

62. Yin X, Wang P, Yang T, Li G, Teng X, Huang W, et al. Identification of key modules and genes associated with breast cancer prognosis using WGCNA and ceRNA network analysis. Aging (Albany NY). 2020;13(2):2519–38.

63. Niemira M, Collin F, Szalkowska A, Bielska A, Chwialkowska K, Reszec J, et al. Molecular Signature of Subtypes of Non-Small-Cell Lung Cancer by Large-Scale Transcriptional Profiling: Identification of Key Modules and Genes by Weighted Gene Co-Expression Network Analysis (WGCNA). Cancers (Basel). 2019;12(1).

64. Chen M, Yan J, Han Q, Luo J, Zhang Q. Identification of hub-methylated differentially expressed genes in patients with gestational diabetes mellitus by multi-omic WGCNA basing epigenome-wide and transcriptome-wide profiling. J Cell Biochem. 2020;121(5-6):3173–84.

65. Huang Z, Ma A, Yang S, Liu X, Zhao T, Zhang J, et al. Transcriptome analysis and weighted gene co-expression network reveals potential genes responses to heat stress in turbot Scophthalmus maximus. Comp Biochem Physiol Part D Genomics Proteomics. 2020;33:100632.

66. Zhang L, Zhang Q, Li W, Zhang S, Xi W. Identification of key genes and regulators associated with carotenoid metabolism in apricot (Prunus armeniaca) fruit using weighted gene coexpression network analysis. BMC Genomics. 2019;20(1):876.

67. Traylor-Knowles N, Connelly MT, Young BD, Eaton K, Muller EM, Paul VJ, et al. Gene Expression Response to Stony Coral Tissue Loss Disease Transmission in M. cavernosa and O. faveolata From Florida. Frontiers in Marine Science. 2021;8.

68. Baris TZ, Wagner DN, Dayan DI, Du X, Blier PU, Pichaud N, et al. Evolved genetic and phenotypic differences due to mitochondrial-nuclear interactions. Plos Genet. 2017;13(3):e1006517.

69. Campbell-Staton SC, Velotta JP, Winchell KM. Selection on adaptive and maladaptive gene expression plasticity during thermal adaptation to urban heat islands. Nature Communications. 2021;12(1):6195.

70. Healy TM, Brennan RS, Whitehead A, Schulte PM. Tolerance traits related to climate change resilience are independent and polygenic. Global Change Biol. 2018;24(11):5348–60.

71. Brown JH, Gillooly JF, Allen AP, Savage VM, West GB. Toward a Metabolic Theory of Ecology. Ecology. 2004;85(7):1771–89.

72. Killen SS, Glazier DS, Rezende EL, Clark TD, Atkinson D, Willener AS, et al. Ecological Influences and Morphological Correlates of Resting and Maximal Metabolic Rates across Teleost Fish Species. Am Nat. 2016;187(5):592–606.

73. Okie JG, Smith VH, Martin-Cereceda M. Major evolutionary transitions of life, metabolic scaling and the number and size of mitochondria and chloroplasts. Proc Biol Sci. 2016;283(1831).

74. Portner HO, Schulte PM, Wood CM, Schiemer F. Niche dimensions in fishes: an integrative view. Physiol Biochem Zool. 2010;83(5):808–26.

75. Wallace DC. A mitochondrial paradigm of metabolic and degenerative diseases, aging, and cancer: a dawn for evolutionary medicine. Annu Rev Genet. 2005;39:359–407.

76. Pettersen AK, Marshall DJ, White CR. Understanding variation in metabolic rate. Journal of Experimental Biology. 2018;221(1).

77. Schulte PM. The effects of temperature on aerobic metabolism: towards a mechanistic understanding of the responses of ectotherms to a changing environment. J Exp Biol. 2015;218(Pt 12):1856–66.

78. Clark TD, Jeffries KM, Hinch SG, Farrell AP. Exceptional aerobic scope and cardiovascular performance of pink salmon (Oncorhynchus gorbuscha) may underlie resilience in a warming climate. J Exp Biol. 2011;214(Pt 18):3074–81.

79. Clark TD, Ryan T, Ingram BA, Woakes AJ, Butler PJ, Frappell PB. Factorial aerobic scope is independent of temperature and primarily modulated by heart rate in exercising Murray cod (Maccullochella peelii peelii). Physiol Biochem Zool. 2005;78(3):347–55.

80. Jensen DL, Overgaard J, Wang T, Gesser H, Malte H. Temperature effects on aerobic scope and cardiac performance of European perch (Perca fluviatilis). J Therm Biol. 2017;68(Pt B):162–9.

81. Nyboer EA, Chapman LJ. Cardiac plasticity influences aerobic performance and thermal tolerance in a tropical, freshwater fish at elevated temperatures. J Exp Biol. 2018;221(Pt 15).

82. Chung DJ, Bryant HJ, Schulte PM. Thermal acclimation and subspecies-specific effects on heart and brain mitochondrial performance in a eurythermal teleost (Fundulus heteroclitus). Journal of Experimental Biology. 2017;220(8):1459–71.

83. Gould SJ. The Panda’s Thumb. New York/London: W. W. Norton & Company, Inc; 1992.

84. Gould SJ, Lewontin RC. The spandrels of San Marco and the Panglossian paradigm: a critique of the adaptationist programme. Proc R Soc Lond B Biol Sci. 1979;205(1161):581–98.

85. Linquist S, Doolittle WF, Palazzo AF. Getting clear about the F-word in genomics. Plos Genet. 2020;16(4):e1008702.

86. Becker CD, Genoway RG. Evaluation of the critical thermal maximum for determining thermal tolerance of freshwater fish. Environmental Biology of Fishes. 1979;4:245–56.

87. Picelli S, Bjorklund AK, Reinius B, Sagasser S, Winberg G, Sandberg R. Tn5 transposase and tagmentation procedures for massively scaled sequencing projects. Genome Res. 2014;24(12):2033–40.

88. Anders S, Huber W. Differential expression analysis for sequence count data. Genome Biol. 2010;11(10):R106.

89. Huerta-Cepas J, Forslund K, Coelho LP, Szklarczyk D, Jensen LJ, von Mering C, et al. Fast Genome-Wide Functional Annotation through Orthology Assignment by eggNOG-Mapper. Mol Biol Evol. 2017;34(8):2115–22.

90. Wu T, Hu E, Xu S, Chen M, Guo P, Dai Z, et al. clusterProfiler 4.0: A universal enrichment tool for interpreting omics data. Innovation (N Y). 2021;2(3):100141.

91. Kanehisa M, Sato Y. KEGG Mapper for inferring cellular functions from protein sequences. Protein Sci. 2020;29(1):28–35.

92. Kanehisa M, Sato Y, Kawashima M, Furumichi M, Tanabe M. KEGG as a reference resource for gene and protein annotation. Nucleic Acids Res. 2016;44(D1):D457–62.

93. Maere S, Heymans K, Kuiper M. BiNGO: a Cytoscape plugin to assess overrepresentation of gene ontology categories in biological networks. Bioinformatics. 2005;21(16):3448–9.

